# Evolutionarily ancient role of cholecystokinin-type neuropeptide signalling as an inhibitory regulator of feeding-related processes revealed in an echinoderm

**DOI:** 10.1101/2020.12.11.417543

**Authors:** Ana B. Tinoco, Antón Barreiro-Iglesias, Luis Alfonso Yañez-Guerra, Jérôme Delroisse, Ya Zhang, Elizabeth F. Gunner, Cleidiane Zampronio, Alexandra M. Jones, Michaela Egertová, Maurice R. Elphick

**Affiliations:** Queen Mary University of London, School of Biological & Chemical Sciences, Mile End Road, London, E1 4NS, UK; School of Life Sciences and Proteomics, Research Technology Platform, University of Warwick, Coventry, UK; Department of Functional Biology, CIBUS, Faculty of Biology, Universidade de Santiago de Compostela, 15782, Santiago de Compostela, Spain; Living Systems Institute, University of Exeter, Exeter EX4 4QD, UK; University of Mons, Biology of Marine Organisms and Biomimetics Unit, 7000 Mons, Belgium

## Abstract

Cholecystokinin (CCK) / sulfakinin (SK)-type neuropeptides regulate feeding and digestion in chordates and protostomes (e.g. insects). Here we characterised CCK/SK-type signalling for the first time in a non-chordate deuterostome - the starfish *Asterias rubens* (phylum Echinodermata). In this species, two neuropeptides (ArCCK1, ArCCK2) derived from the precursor protein ArCCKP act as ligands for a CCK/SK-type receptor (ArCCKR) and are expressed in the nervous system, digestive system, tube feet and body wall. Furthermore, ArCCK1 and ArCCK2 cause dose-dependent contraction of cardiac stomach, tube foot and body wall apical muscle preparations *in vitro* and injection of these neuropeptides *in vivo* triggers cardiac stomach retraction and inhibition of the onset of feeding in *A. rubens*. Thus, an evolutionarily ancient role of CCK/SK-type neuropeptides as inhibitory regulators of feeding-related processes in the Bilateria has been conserved in the unusual and unique context of the extra-oral feeding behaviour and pentaradial body plan of an echinoderm.

## Introduction

The peptide hormones cholecystokinin (CCK) and gastrin were discovered and named on account of their effects as stimulators of gall bladder contraction and gastric secretion of pepsin/acid, respectively, in mammals (Edkins 1906, Ivy 1929). Determination of the structures of CCK and gastrin revealed that they have the same C-terminal structural motif (Trp-Met-Asp-Phe-NH_2_), indicative of a common evolutionary origin (Gregory et al. 1964, Mutt and Jorpes 1968). However, CCK and gastrin are derived from different precursor proteins, which are subject to cell/tissue-specific processing to give rise to bioactive peptides of varying length (e.g. CCK-8 and CCK-33; gastrin-17 and gastrin-34) (Boel et al. 1983, Deschenes et al. 1984, Rehfeld et al. 2007). Furthermore, CCK and gastrin have a tyrosine residue at positions seven and six from the C-terminal amide, respectively, which can be sulphated post-translationally (Rehfeld et al. 2007). The effects of CCK and gastrin in mammals are mediated by two G-protein coupled receptors (GPCRs), CCK-A (CCKR1) and CCK-B (CCKR2), with both sulphated and non-sulphated forms of CCK and gastrin acting as ligands for the CCK-B receptor, whilst the CCK-A receptor is selectively activated by sulphated CCK (Deweerth et al. 1993, Dufresne et al. 2006, Kopin et al. 1992, Lee et al. 1993, Noble and Roques 1999, Wank et al. 1992). Mediated by these receptors, gastrin and CCK have a variety of physiological/behavioural effects in mammals. Thus, in the gastrointestinal system gastrin stimulates growth of the stomach lining, gastric contractions and gastric emptying (Crean et al. 1969, Dockray et al. 2005, Gregory and Tracy 1964, Vizi et al. 1973), whilst CCK stimulates pancreatic enzyme secretion, contraction of the pyloric sphincter and intestinal motility (Chen et al. 2004, Gutiérrez et al. 1974, Harper and Raper 1943, Rehfeld 2017, Shaw and Jones 1978, Vizi et al. 1973). Furthermore, CCK also has behavioural effects that include inhibition of food intake as a mediator of satiety and stimulation of aggression and anxiogenesis (Chandra and Liddle 2007, Gibbs et al. 1973, Singh et al. 1991, Smith et al. 1981).

Phylogenomic studies indicate that genome duplication in a common ancestor of the vertebrates gave rise to genes encoding CCK-type and gastrin-type precursor proteins (Dupre and Tostivint 2014). Accordingly, invertebrate chordates that are the closest extant relatives of vertebrates (e.g. the urochordate *Ciona intestinalis*) have a single gene encoding a “hybrid” CCK/gastrin-like peptide (e.g. cionin) with a sulphated tyrosine residue at both positions six and seven from the C-terminal amide (Johnsen and Rehfeld 1990, Monstein et al. 1993, Thorndyke and Dockray 1986). Furthermore, CCK-type peptides stimulate gastric enzyme secretion in the sea–squirt *Styela clava*, providing evidence of evolutionarily ancient roles as regulators of gastrointestinal physiology in chordates (Bevis and Thorndyke 1981, Thorndyke and Bevis 1984).

Evidence that the phylogenetic distribution of CCK/gastrin-type peptides may extend beyond chordates to other phyla was first obtained with the detection of substances immunoreactive with antibodies to CCK and/or gastrin in a variety of invertebrates, including arthropods, annelids, molluscs and cnidarians (Dockray et al. 1981, El-Salhy et al. 1980, Grimmelikhuijzen et al. 1980, Kramer et al. 1977, Larson and Vigna 1983, Rzasa et al. 1982). However, molecular evidence of the evolutionary antiquity of CCK/gastrin-type signalling was obtained with the purification and sequencing of a CCK-like peptide named leucosulfakinin, which was isolated from the insect (cockroach) *Leucophaea maderae* (Nachman et al. 1986a). Subsequently, GPCRs that are homologs of the vertebrate CCKA/CCKB-type receptors have been identified and pharmacologically characterised as receptors for sulfakinin (SK)-type peptides in a variety of insects, including *Drosophila melanogaster* (Bloom et al. 2019, Kubiak et al. 2002, Yu et al. 2013b, Yu and Smagghe 2014b). Furthermore, investigation of the physiological roles of SK-type signalling in insects has revealed similarities with findings from vertebrates. Thus, in several insect species SK-type peptides have myotropic effects on the gut (Al-Alkawi et al. 2017, Marciniak et al. 2011, Nachman et al. 1986a, Nachman et al. 1986b, Nichols 2007, Palmer et al. 2007, Predel et al. 2001, Schoofs et al. 1990) and/or affect digestive enzyme release (Harshini et al. 2002a, b, Nachman et al. 1997, Zels et al. 2015). Furthermore, at a behavioural level there is evidence that SK-type peptides act as satiety factors (Al-Alkawi et al. 2017, Bloom et al. 2019, Downer et al. 2007, Maestro et al. 2001, Meyering-Vos and Muller 2007, Nässel and Zandawala 2019, Nichols et al. 2008, Wei et al. 2000, Yu et al. 2013a, Yu et al. 2013b, Yu and Smagghe 2014b, Zels et al. 2015) and regulate locomotion and aggression in insects (Chen et al. 2012, Nässel and Williams 2014, Nässel and Zandawala 2019, Nichols et al. 2008).

The discovery and functional characterisation of SK-type signalling in insects and other arthropods indicated that the evolutionary origin CCK/SK-type signalling can be traced back to the common ancestor of the Bilateria. Consistent with this hypothesis, CCK/SK-type signalling systems have been discovered in a variety of protostome invertebrates, including the nematode *Caenorhabditis elegans*, the mollusc *Crassostrea gigas* and the annelid *Capitella teleta* (Janssen et al. 2008, Mirabeau and Joly 2013, Schwartz et al. 2018). Furthermore, some insights into the physiological roles of CCK/SK-type signalling in non-arthropod protostomes have been obtained, including causing a decrease in the frequency of spontaneous contractions of the *C. gigas* hindgut (Schwartz et al. 2018), stimulation of digestive enzyme secretion in *C. elegans* and *Pecten maximus* (Janssen et al. 2008, Nachman et al. 1997) and evidence of a role in regulation of feeding and energy storage in *C. gigas* (Schwartz et al. 2018).

Little is known about CCK-type signalling in the Ambulacraria (echinoderms and hemichordates) - deuterostome invertebrates that occupy an ‘intermediate’ phylogenetic position with respect to chordates and protostomes (Furlong and Holland 2002, Telford et al. 2015). Prior to the genome sequencing era, use of immunohistochemical methods revealed CCK-like immunoreactive cells in the intestine of sea cucumbers (Phylum Echinodermata) and vertebrate CCK/gastrin-type peptides were found to cause relaxation of sea cucumber intestine (García-Arrarás et al. 1991). More recently, analysis of transcriptome/genome sequence data has enabled identification of transcripts/genes encoding CCK-type peptide precursors and CCK-type receptors in echinoderms and hemichordates (Burke et al. 2006, Chen et al. 2019, Jekely 2013, Mirabeau and Joly 2013, Semmens et al. 2016, Zandawala et al. 2017). However, functional characterisation of native CCK-type peptides and receptors has yet to be reported for an echinoderm or hemichordate species. We have established the common European starfish *Asterias rubens* as an experimental model for molecular and functional characterisation of neuropeptides, obtaining novel insights into the evolution and comparative physiology of several neuropeptide signalling systems (Cai et al. 2018, Elphick et al. 2018, Lin et al. 2017a, Lin et al. 2018, Odekunle et al. 2019, Semmens and Elphick 2017, Tian et al. 2017, Tian et al. 2016, Tinoco et al. 2018, Yáñez-Guerra et al. 2018, Yáñez-Guerra et al. 2020, Zhang et al. 2020). Accordingly, here we used *A. rubens* to enable the first detailed molecular, anatomical and pharmacological analysis of CCK-type signalling in an echinoderm.

## Results

### Cloning and sequencing of a cDNA encoding ArCCKP

Analysis of *A. rubens* neural transcriptome sequence data has revealed the presence of a CCK-type precursor in *A. rubens*, which was named ArCCKP (Semmens et al. 2016). Here, cloning and sequencing of a cDNA encoding ArCCKP confirmed the sequence obtained from transcriptome data (Figure 1 – figure supplement 1).

### Identification of ArCCKP-derived neuropeptides in extracts of *A. rubens* radial nerve cords

ArCCKP comprises two putative CCK-like neuropeptide sequences that are bounded by dibasic or tetrabasic cleavage sites. Both neuropeptide sequences have a C-terminal glycine residue, which is a potential substrate for post-translational amidation, and both neuropeptide sequences contain a tyrosine residue, which could be either sulphated or non-sulphated (ns) in the mature neuropeptides. Furthermore, an N-terminal glutamine residue (Q) in one of the neuropeptide sequences is a potential substrate for N-terminal pyroglutamylation (pQ) (Figure 1 – figure supplement 1; Figure 1 - figure supplement 2a).

LC-MS-MS analysis of *A. rubens* radial nerve cord extracts revealed the presence of four CCK-type peptides derived from ArCCKP: pQSKVDDY(SO_3_H)GHGLFW-NH_2_ (ArCCK1; Figure 1 – figure supplement 2b), pQSKVDDYGHGLFW-NH_2_ [ArCCK1(ns); Figure 1 – figure supplement 2c], GGDDQY(SO_3_H)GFGLFF-NH_2_ (ArCCK2; Figure 1 – figure supplement 2d) and GGDDQYGFGLFF-NH_2_ [ArCCK2(ns); Figure 1 – figure supplement 2e]. Thus, mass spectrometry confirmed that i). the peptides are C-terminally amidated, ii). the peptides are detected with or without tyrosine sulphation and iii). an N-terminal glutamine is post-translationally converted to pyroglutamate in the mature ArCCK1 and ArCCK1(ns) peptides.

Having determined the structures of CCK-type neuropeptides derived from ArCCKP, the sequences of ArCCK1 and ArCCK2 were aligned with the sequences of CCK-type peptides that have been identified in other taxa (Figure 1). This revealed a number of evolutionarily conserved features, including a tyrosine residue (typically sulphated) and a C-terminal amide group that are separated by five to seven intervening residues. The C-terminal residue in the majority of CCK-type peptides, including ArCCK2, is a phenylalanine residue. However, ArCCK1 is atypical in having a C-terminal tryptophan residue, which is also a feature of a CCK-type peptide in the bivalve mollusc *C. gigas*.

**Figure 1.**
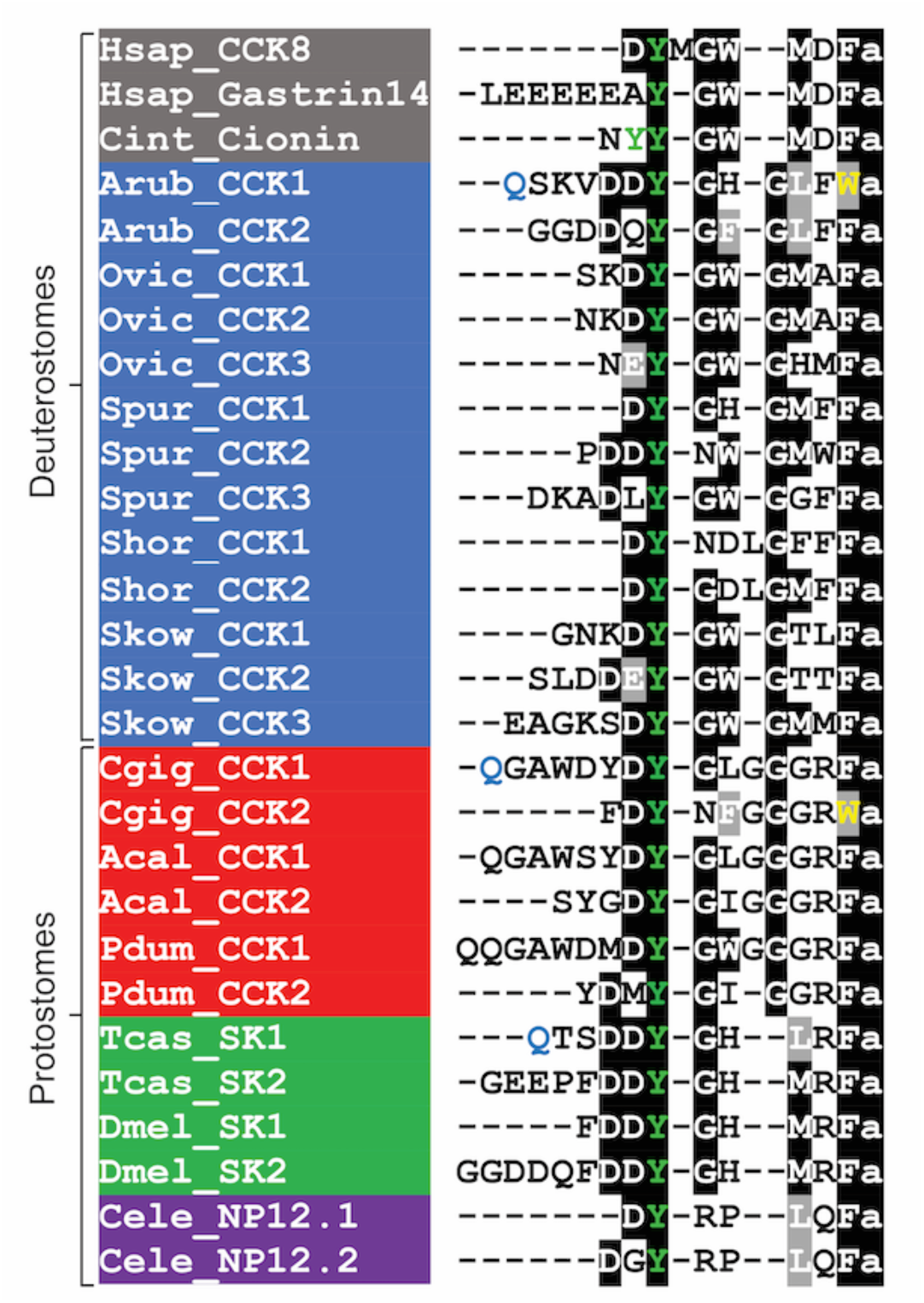
Comparison of the *A. rubens* CCK-type neuropeptides ArCCK1 and ArCCK2 with CCK/SK-type neuropeptides from other taxa. Conserved residues are highlighted, with conservation in more than 70% of sequences highlighted in black and conservative substitutions highlighted in grey. Experimentally verified conversion of an N-terminal glutamine residue (Q) to pyroglutamate in the mature peptide is indicated by the letter Q being shown in light blue. Tyrosine (Y) residues that are known or predicted to be subject to post-translational sulphation are shown in green. The C-terminal tryptophan (W) in ArCCK1 and in a *C. gigas* CCK-type peptide are shown in yellow to highlight that this feature is atypical of CCK-type peptides. Predicted or experimentally verified C-terminal amides are shown as the letter “a” in lowercase. Species names are highlighted in taxon-specific colours: grey (Chordata), blue (Ambulacraria), red (Lophotrochozoa), green (Arthropoda) and purple (Nematoda). Abbreviations are as follows: Acal (*Aplysia californica*), Arub (*Asterias rubens*), Cele (*Caenorhabditis elegans*), Cgig (*Crassostrea gigas*), Cint (*Ciona intestinalis*), Dmel (*Drosophila melanogaster*), Hsap (*Homo sapiens*), Ovic (*Ophionotus victoriae*), Pdum (*Platynereis dumerilii*), Shor (*Stichopus horrens*), Skow (*Saccoglossus kowalevskii*), Spur (*Strongylocentrotus purpuratus*), Tcas (*Tribolium castaneum*). The accession numbers of the sequences are listed in Figure 1 – source data 1. The nucleotide sequence of a cloned cDNA encoding the *A. rubens* cholecystokinin-type precursor is shown in Figure 1 – figure supplement 1. Mass spectroscopic analysis of the structures of the peptides derived from the *A. rubens* cholecystokinin-type precursor is presented in Figure 1 – figure supplement 2. The raw data for the results shown in Figure 1 – figure supplement 2 can be found in **Figure 1 – source data 2**.

### Identification of a CCK-type receptor in *A. rubens*

BLAST analysis of *A. rubens* neural transcriptome sequence data identified a transcript that encodes a 434-residue protein (ArCCKR) that shares high sequence similarity with CCK-type receptors from other taxa (Figure 2 – figure supplement 1). Phylogenetic analysis revealed that ArCCKR groups within a clade including CCK-type receptors that have been pharmacologically characterised in other taxa, including the human and mouse CCK/gastrin receptors CCKR1 and CCKR2, the *C. intestinalis* cionin receptors CioR1 and CioR2, the *Drosophila melanogaster* sulfakinin receptors SKR1 and SKR2, and the recently characterised *C. gigas* receptors CCKR1 and CCKR2 (Figure 2). Thus, this demonstrates that ArCCKR is an ortholog of CCK/SK-type receptors that have been characterised in other taxa. Furthermore, reflecting known animal phylogenetic relationships, ArCCKR is positioned within a branch of the tree that comprises CCK-type receptors from deuterostomes, and more specifically it is positioned within an ambulacrarian clade that comprises CCK-type receptors from other echinoderms and from the hemichordate *Saccoglossus kowalevskii*.

**Figure 2.**
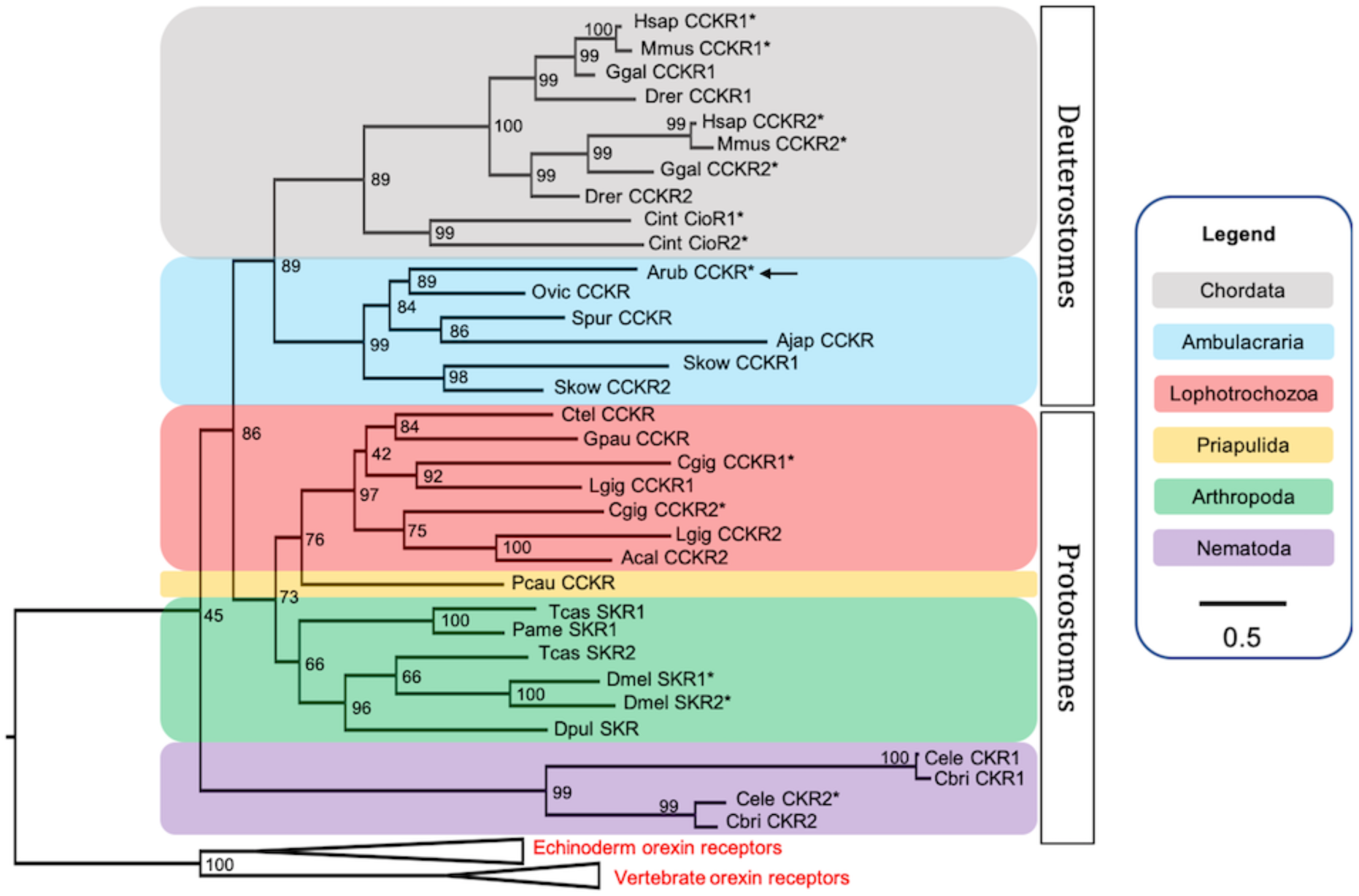
Phylogenetic tree showing that the predicted *A. rubens* CCK-type receptor (ArCCKR; arrow) is an ortholog of CCK-type receptors in other taxa, which include receptors (*) for which the peptide ligands have been identified experimentally. The tree was generated using the maximum likelihood method (Guindon et al. 2009), with the percentage of replicate trees in which two or more sequences form a clade in a bootstrap test (1000 replicates) shown at the node of each clade (Zhaxybayeva and Gogarten 2002). Orexin receptors were included as an outgroup. The tree is drawn to scale, with branch lengths in the same units as those of the evolutionary distances used to infer the phylogenetic tree. Evolutionary analyses were conducted using the IQ-tree server (Trifinopoulos et al. 2016). Taxa are colour-coded as explained in the key. Abbreviations of species names are as follows: Acal (*Aplysia californica*), Ajap (*Apostichopus japonicus*), Arub (*Asterias rubens*), Cbri (*Caenorhabditis briggsae*), Cele (*Caenorhabditis elegans*), Cgig (*Crassostrea gigas*), Cint (*Ciona intestinalis*), Ctel (*Capitella teleta*), Dmel (*Drosophila melanogaster*), Dpul (*Daphnia pulex*), Drer (*Danio rerio*), Ggal (*Gallus gallus*), Gpau (*Glossoscolex paulistus*), Hsap (*Homo sapiens*), Lgig (*Lottia gigantea*), Mmus (*Mus musculus*), Ovic (*Ophionotus victoriae*), Pame (*Periplaneta americana*), Pcau (*Priapulus caudatus*), Skow (*Saccoglossus kowalevskii*), Spur (*Strongylocentrotus purpuratus*), Tcas (*Tribolium castaneum*). The accession numbers of the sequences used for this phylogenetic tree are listed in Figure 2 – source data 1. The nucleotide sequence and the predicted topology of ArCCKR are shown in Figure 2 – figure supplement 1 and Figure 2 – figure supplement 2, respectively.

Analysis of the amino acid sequence of ArCCKR using the Protter tool (Omasits et al. 2014) revealed seven predicted transmembrane domains, as expected for a GPCR, and three potential N-glycosylation sites in the predicted extracellular N-terminal region of the receptor (Figure 2 – figure supplement 2).

### ArCCK1 and ArCCK2 are ligands for ArCCKR

Previous studies on other species have revealed that sulphation of the tyrosine residue in CCK-type peptides is often important for receptor activation and bioactivity (Dufresne et al. 2006, Kubiak et al. 2002, Schwartz et al. 2018, Sekiguchi et al. 2012, Yu et al. 2015). Accordingly, neuropeptides derived from ArCCKP were detected in *A. rubens* radial nerve cord extracts with sulphated tyrosines (ArCCK1 and ArCCK2; Figure 1 – figure supplement 2b, d). Therefore, the sulphated peptides ArCCK1 and ArCCK2 were synthesized and tested as ligands for ArCCKR. However, because non-sulphated forms of ArCCK1 (ArCCK1(ns)) and ArCCK2 (ArCCK(ns)) were also detected *A. rubens* radial nerve extracts (Figure 1 – figure supplement 2c, e), we also synthesized and tested ArCCK2(ns) to investigate if absence of tyrosine sulphation affects receptor activation. Using CHO-K1 cells expressing aequorin as an assay system, ArCCK1, ArCCK2 and ArCCK2(ns) did not elicit luminescence responses when tested on cells transfected with an empty vector (Figure 3). However, all three peptides caused concentration-dependent stimulation of luminescence in CHO-K1 cells transfected with ArCCKR (Figure 3). The EC_50_ values for ArCCK1, ArCCK2 and ArCCK2(ns) were 0.25 nM (Figure 3a), 0.12 nM (Figure 3b) and 48 µM (Figure 3c), respectively. Thus, although activation of ArCCKR was observed *in vitro* with ArCCK2(ns) (Figure 3c), this peptide is five to six orders of magnitude less potent than ArCCK1 and ArCCK2 as a ligand for ArCCKR. This indicates that the non-sulphated peptides ArCCK1(ns) and ArCCK2(ns) that were detected in *A. rubens* radial nerve extracts are unlikely to have physiological effects *in vivo*. Furthermore, two other *A. rubens* neuropeptides that share modest C-terminal sequence similarity with the *A. rubens* CCK-type peptides - the SALMFamide neuropeptide S2 (SGPYSFNSGLTF-NH_2_) and the tachykinin-like peptide ArTK2 (GGGVPHVFQSGGIF-NH_2_) - were found to be inactive when tested as ligands for ArCCKR at concentrations ranging from 10^-12^ to 10^-4^ M) (Figure 3 – figure supplement 1), demonstrating the specificity of ArCCK1 and ArCCK2 as ligands for ArCCKR.

**Figure 3.**
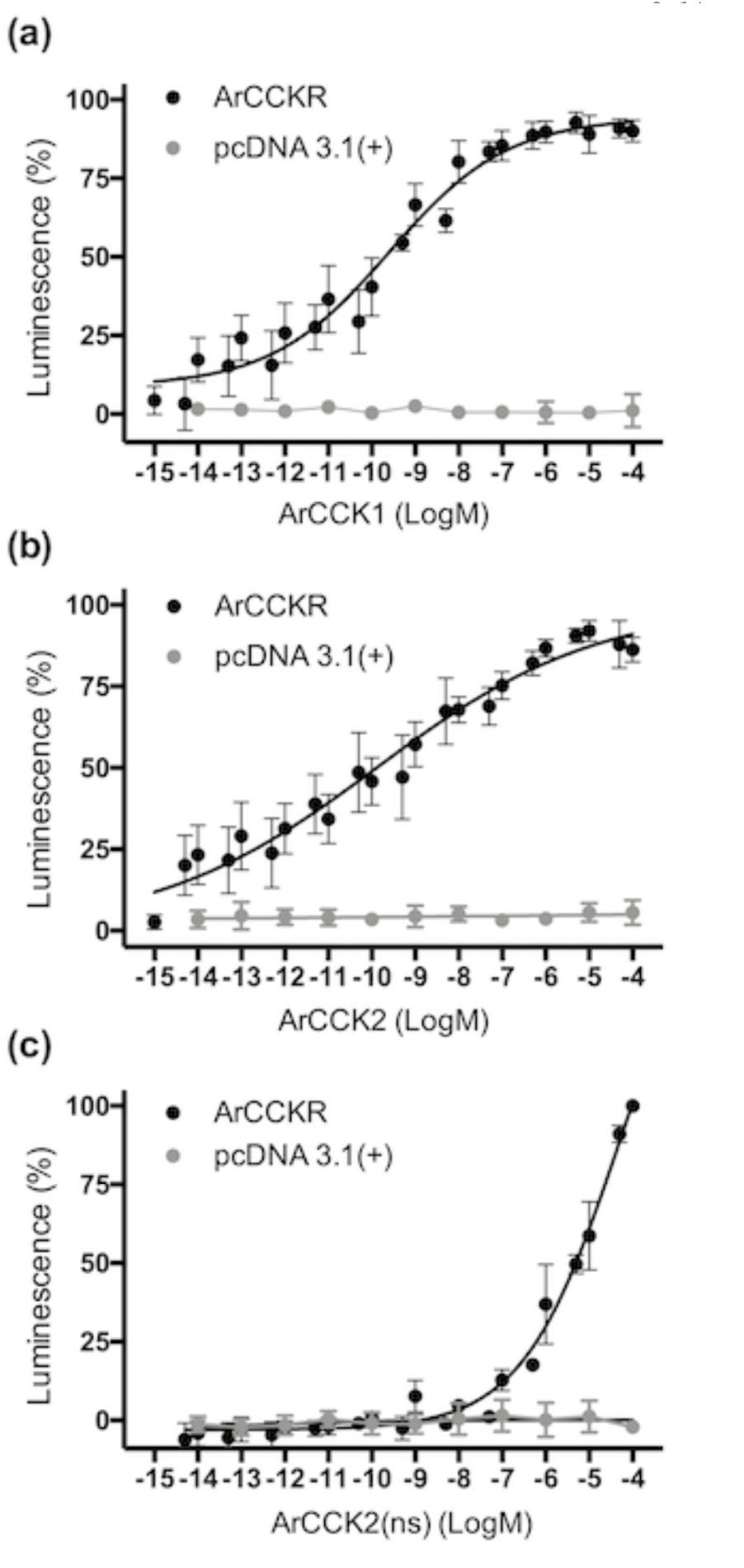
Experimental demonstration that the *A. rubens* CCK-type peptides ArCCK1 and ArCCK2 act as ligands for the *A. rubens* CCK-type receptor ArCCKR. The sulphated peptides ArCCK1 **(a),** ArCCK2 **(b)** and the non-sulphated peptide ArCCK2(ns) **(c)** trigger dose-dependent luminescence in CHO-K1 cells stably expressing mitochondrial targeted apoaequorin (G5A) that were co-transfected with plasmids encoding the promiscuous human G-protein Gα16 and ArCCKR (black). Control experiments where cells were transfected with an empty pcDNA 3.1(+) vector are shown in grey. Each point represents mean values (± s.e.m) from at least four independent experiments performed in triplicate. Luminescence is expressed as a percentage of the maximal response observed in each experiment. The EC_50_ values for ArCCK1 (a) and ArCCK2 (b) are 0.25 nM and 0.12 nM, respectively. In comparison, the absence of tyrosine (Y) sulphation in ArCCK2(ns) (c) causes a massive loss of potency (EC_50_ = 48 µM), indicating that the sulphated peptides act as ligands for ArCCKR physiologically. A graph showing the selectivity of ArCCKR as a receptor for CCK-type peptides is presented in Figure 3 – figure supplement 1

### Localisation of ArCCKP expression in *A. rubens* using mRNA *in situ* hybridization

To gain anatomical insights into the physiological roles of CCK-type neuropeptides in starfish, mRNA *in situ* hybridisation methods were employed to enable analysis of the distribution of the ArCCKP transcript in *A. rubens*. As described below and illustrated in Figure 4, expression of ArCCKP was observed in the central nervous system, digestive system, body wall and tube feet.

**Figure 4.**
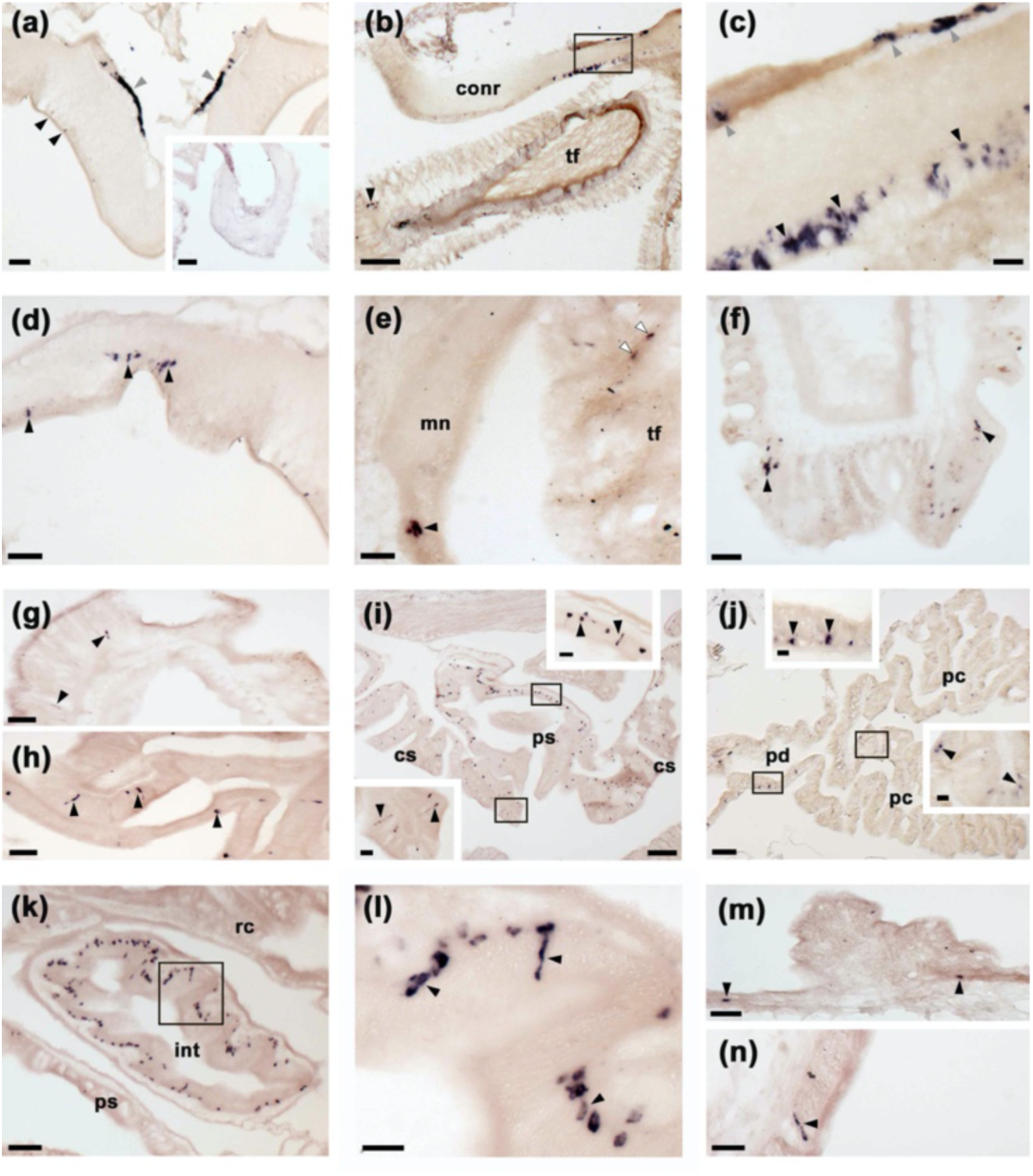
Localisation of ArCCKP expression in *A. rubens* using mRNA *in situ* hybridization. **(a)** Using antisense probes, ArCCKP-expressing cells are revealed in the ectoneural (black arrowheads) and hyponeural (grey arrowheads) regions of a radial nerve cord. The specificity of staining with antisense probes is demonstrated by the absence of staining in a radial nerve cord section incubated with sense probes (see inset). **(b)** ArCCKP-expressing cells in the circumoral nerve ring (see boxed area) and in the disk region of a peri-oral tube foot (black arrowhead). **(c)** High magnification image of the boxed area in **(b)**, showing stained cells in the ectoneural (black arrowheads) and hyponeural (grey arrowheads) regions of the circumoral nerve ring. **(d)** ArCCKP-expressing cells in a lateral branch of the radial nerve cord (black arrowheads). **(e)** ArCCKP-expressing cells adjacent to the marginal nerve (black arrowhead) and in the stem of a tube foot (white arrowheads). **(f)** ArCCKP-expressing cells (black arrowheads) adjacent to the basal nerve ring in the disk region of a tube foot. **(g)** ArCCKP-expressing cells (black arrowheads) in the mucosal layer of the oesophagus. **(h)** ArCCKP-expressing cells (black arrowheads) in the mucosal layer of the cardiac stomach. **(i)** ArCCKP-expressing cells (black arrowheads) in the cardiac stomach and pyloric stomach, with the boxed regions shown at higher magnification in the insets. **(j)** ArCCKP-expressing cells (black arrowheads) in the pyloric duct and pyloric caeca, with the boxed regions shown at higher magnification in the insets. **(k, l)** ArCCKP-expressing cells (black arrowheads) in an oblique section of the intestine; the boxed region in **(k)** is shown at higher magnification in **(l)**. **(m,n)** ArCCKP-expressing cells (black arrowheads) in the external epithelium of the body wall. Abbreviations: conr, circumoral nerve ring; int, intestine; mn, marginal nerve; pc, pyloric caecum; pd, pyloric duct; ps, pyloric stomach; rc, rectal caeca; tf, tube foot. Scale bars: [b), i), j)] = 120 μm; [a), a-inset), k)] = 60 μm; [d), e), f), g), h), m)] = 32 μm; [c), i-insets), j-insets), l), n)] = 16 μm.

The central nervous system of *A. rubens* comprises radial nerve cords that extend along the oral side of each arm, with two rows of tube feet (locomotory organs) on either side. The five radial nerve cords are linked by a circumoral nerve ring in the central disk (Pentreath and Cobb 1972). Analysis of ArCCKP mRNA expression revealed stained cells in both the ectoneural and hyponeural regions of the radial nerve cords (Figure 4a) and circumoral nerve ring (Figure 4b, c). Furthermore, the specificity of staining observed with anti-sense probes was confirmed by an absence of staining in tests with sense probes (Figure 4a). Stained cells were also revealed in the ectoneural segmental branches of the radial nerve cords (Figure 4d) and in the marginal nerves (Figure 4e), which run parallel with the radial nerve cords lateral to the outer row of tube feet. ArCCKP-expressing cells were also revealed in tube feet, with stained cells located in the podium proximal to its junction with the radial and marginal nerves (Figure 4e) and in the tube foot disk (Figure 4f). In the digestive system, ArCCKP-expressing cells were revealed in the mucosa of the oesophagus (Figure 4g), cardiac stomach (Figure 4i, h), pyloric stomach (Figure 4i), pyloric ducts (Figure 4j), pyloric caeca (Figure 4j) and intestine (Figure 4k, l). ArCCKP expressing cells were also revealed in the external epithelium of the body wall (Figure 4m, n).

### Immunohistochemical localisation of ArCCK1 in *A. rubens*

Use of mRNA *in situ* hybridisation (see above) revealed the location of cells expressing ArCCKP in *A. rubens*. However, a limitation of this technique is that it does not reveal the axonal processes of neuropeptidergic neurons. Therefore, to enable this using immunohistochemistry, we generated and affinity-purified rabbit antibodies to ArCCK1. ELISA analysis of antiserum revealed the presence of antibodies to the ArCCK1 peptide antigen (Figure 5 - figure supplement 1a) and ELISA analysis of affinity-purified antibodies to the ArCCK1 antigen peptide revealed the specificity of these antibodies for ArCCK1 because they do not cross-react with ArCCK2, ArCCK2(ns) or the starfish luqin-type neuropeptide ArLQ (Figure 5 - figure supplement 1b). Immunohistochemical tests with affinity-purified ArCCK1 antibodies revealed extensive immunostaining in sections of *A. rubens*, as described in detail below and illustrated in Figure 5.

**Figure 5.**
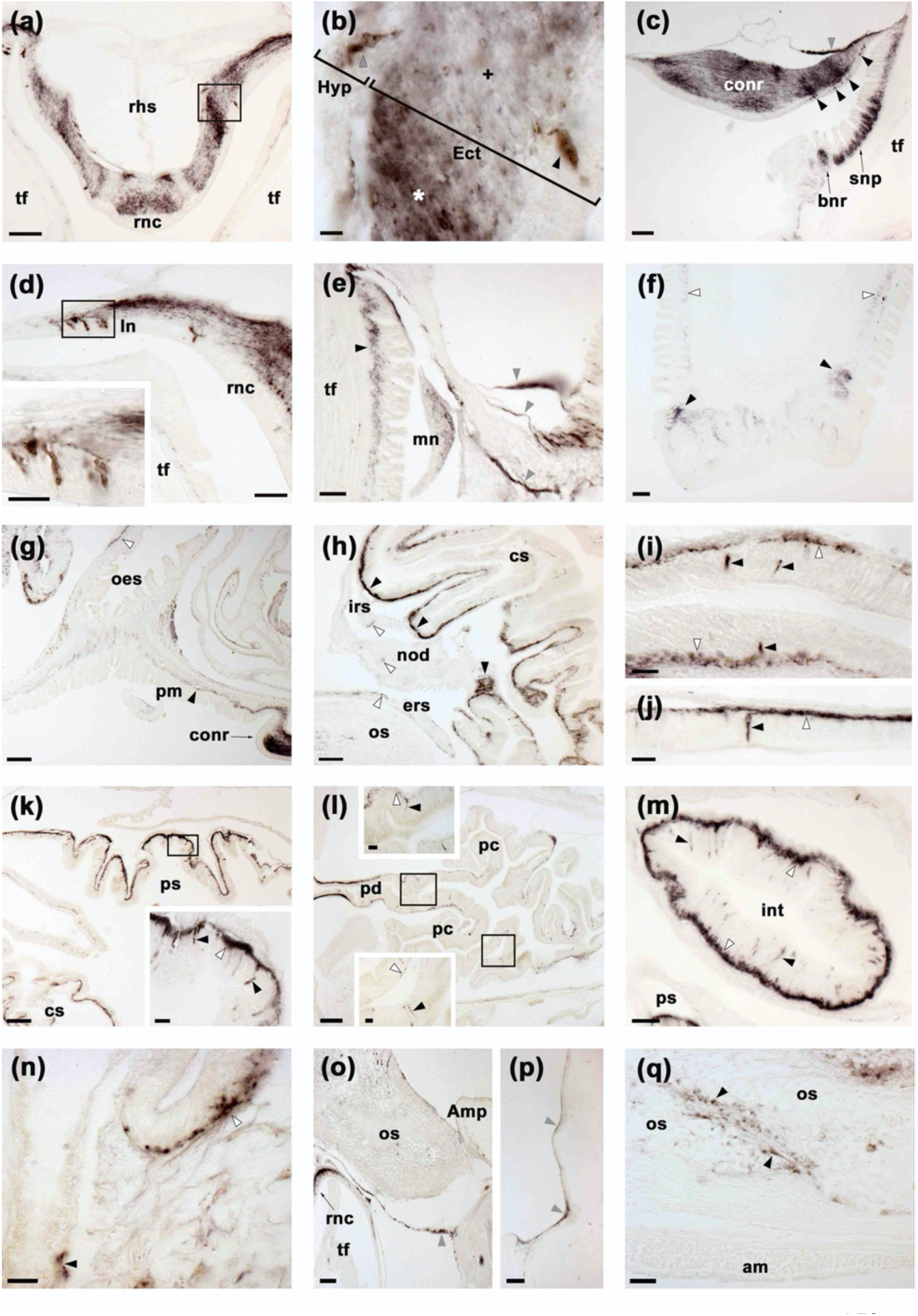
Localisation of ArCCK1 expression in *A. rubens* using immunohistochemistry. **(a)** ArCCK1-immunoreactivity (ArCCK1-ir) in a transverse section of the V-shaped radial nerve cord, with bilaterally symmetrical regional variation in the density of immunostaining in the ectoneural neuropile. **(b)** High magnification image of the boxed region in **(a)**, showing stained cell bodies in the hyponeural (grey arrowhead) and ectoneural (black arrowhead) regions of the radial nerve cord. Regions of the ectoneural neuropile containing a higher (*) and lower (+) densities of immunostained fibres can be seen here. **(c)** ArCCK1-ir in the circumoral nerve ring, with stained cells present in the hyponeural region (grey arrowheads) and in the ectoneural epithelium (black arrowheads); in the ectoneural neuropile there is regional variation in the density of immunostained fibres. Immunostaining can also be seen here in the sub-epithelial nerve plexus and basal nerve ring of an adjacent peri-oral tube foot. **(d)** ArCCK1-ir in cells and fibres in a lateral branch of the radial nerve cord; the inset shows immunostained cells in the boxed region at higher magnification. **(e)** ArCCK1-ir in the marginal nerve, the sub-epithelial nerve plexus of an adjacent tube foot (black arrowhead) and in branches of the lateral motor nerve (grey arrowheads). **(f)** ArCCK1-ir in the sub-epithelial nerve plexus (white arrowheads) and basal nerve ring (black arrowheads) of a tube foot. **(g)** ArCCK1-ir in the basi-epithelial nerve plexus of the peristomial membrane (black arrowhead) and the oesophagus (white arrowhead); immunostaining in the circumoral nerve can also be seen here. **(h)** ArCCK1-ir in the lateral pouches of the cardiac stomach; note that the density of immunostained fibres is highest (black arrowheads) in regions of the mucosa adjacent to the intrinsic retractor strand; immunostaining in the intrinsic retractor strand, nodule and extrinsic retractor strand can also be seen here (white arrowheads). **(i, j)** High magnification images of cardiac stomach tissue showing ArCCK1-ir in cell bodies (black arrowheads) and their processes in the basiepithelial nerve plexus (white arrowheads); note that in **(j)** a process emanating from an immunostained cell body can be seen projecting into the plexus. **(k)** ArCCK1-ir in the cardiac stomach and pyloric stomach; the boxed region is shown at higher magnification in the inset, showing immunostaining in cells (black arrowheads) and the basi-epithelial nerve plexus (white arrowhead). **(l)** ArCCK1-ir in the pyloric duct and pyloric caeca; the boxed regions are shown at higher magnification in the insets, where immunostained cells (black arrowheads) and fibres (white arrowheads) can be seen. **(m)** ArCCK1-ir in an oblique section of the intestine, with immunostained cells in the mucosa (black arrowheads) and intense immunostaining in the basi-epithelial nerve plexus (white arrowheads). **(n)** ArCCK1-ir in the basi-epithelial nerve plexus of the body wall external epithelium (white arrowhead) and in the lining of a papula (black arrowhead). (**o)** ArCCK1-ir in nerve fibres projecting around the base of a tube foot at its junction with the neck of its ampulla. **(p)** ArCCK1-ir in nerve fibres located in the coelomic lining of the lateral region of the body wall. **(q)** ArCCK1-ir in inter-ossicular tissue of the body wall. Abbreviations: am, apical muscle; Amp, ampulla; bnr, basal nerve ring; conr, circumoral nerve ring; cs, cardiac stomach; Ect, ectoneural; ers, extrinsic retractor strand; Hyp, hyponeural; int, intestine; irs, intrinsic retractor strand; ln, lateral nerve; mn, marginal nerve; nod, nodule; oes, oesophagus; os, ossicle; pm, peristomial membrane; pc, pyloric caecum; pd, pyloric duct; ps, pyloric stomach; rhs, radial hemal strand; rnc, radial nerve cord; snp, sub-epithelial nerve plexus; tf, tube foot. Scale bars: [g), k), l)] = 120 μm; [a), c), f), h), o), p)] = 60 μm; [d), e), m), q)] = 32 μm; [d-inset), i), j), k-inset), l-insets), n)] = 16 μm; [b)] = 6 μm. Graphs showing ELISA-based characterisation of the antibodies to ArCCK1 used here for immunohistochemistry are presented in Figure 5 – figure supplement 1.

ArCCK1-immunoreactive (ir) cells were revealed in the ectoneural and hyponeural regions of the radial nerve cords (Figure 5a, b). Furthermore, dense networks of immunostained fibres were revealed in the ectoneural neuropile, with bilaterally symmetrical regional variation in the density of immunostaining (Figure 5b). Likewise, ArCCK1-ir cells were revealed in the ectoneural and hyponeural regions of the circumoral nerve ring, also with regional variation in the density of immunostained fibres in the ectoneural neuropile (Figure 5c). Immunostained cells and/or processes were also revealed in the segmental lateral branches of the radial nerve cords (Figure 5d) and in the marginal nerve cords (Figure 5e). Consistent with the expression of ArCCKP/ArCCK1 in the hyponeural region of radial nerve cords, ArCCK1-ir fibres were revealed in the lateral motor nerves (Figure 5e). In tube feet, ArCCK1-ir fibres were revealed in the sub-epithelial nerve plexus of the podium and in the basal nerve ring of the disk region (Figure 5c,f).

ArCCK1-ir cells and/or fibres were revealed in the mucosa and basiepithelial nerve plexus, respectively, of many regions of the digestive system, including the peristomial membrane (Figure 5g), oesophagus (Figure 5g), cardiac stomach (Figure 5h, i, j), pyloric stomach (Figure 5k), pyloric ducts (Figure 5l), pyloric caeca (Figure 5l) and intestine (Figure 5m). Consistent with patterns of ArCCKP transcript expression (see above), regional differences in the abundance of stained cells and fibres were observed. Regions of the digestive system containing denser populations of CCK1-ir cells and/or fibres include the lateral pouches of the cardiac stomach (Figure 5h), the roof of the pyloric stomach (Figure 5k) and the intestine (Figure 5m).

Immunostaining was observed in the sub-epithelial nerve plexus of the external epithelium of the body wall (Figure 5n) and in papulae (Figure 5n), which are protractable appendages that penetrate through the body wall to enable gas exchange between external seawater and the coelomic fluid (Cobb 1978). Immunostained fibres were also observed in branches of the lateral motor nerves located in the coelomic lining of the body wall (Figure 5o,p). However, no staining was observed in the apical muscle - a thickening of longitudinally orientated muscle that is located under the coelomic epithelial layer of the body wall along the aboral midline of each arm (Figure 5q). The bulk of the body wall in *A. rubens* is comprised of calcite ossicles that are interconnected by muscles and collagenous tissue and ArCCK1-ir fibres were revealed in the inter-ossicular tissue (Figure 5q).

### ArCCK1 and ArCCK2 cause concentration-dependent contraction of *in vitro* cardiac stomach, tube foot and apical muscle preparations from *A. rubens*

Informed by the localisation of ArCCKP/ArCCK1 expression in the cardiac stomach and tube feet of *A. rubens*, we tested the effects of ArCCKP-derived neuropeptides on *in vitro* preparations of these organs. Both ArCCK1 and ArCCK2 caused concentration-dependent contraction of cardiac stomach preparations when tested at concentrations ranging from 1 nM to 1 μM (Figure 6a, b). ArCCK2(ns) caused modest contraction of cardiac stomach preparations by comparison with ArCCK2 (Figure 6b), and comparison of the ArCCK2 and ArCCK2(ns) data using a 2-way ANOVA revealed a significant difference in the effects of the peptides on cardiac stomach preparations, irrespective of concentration (P < 0.0001). However, 2-way ANOVA analysis revealed no significant difference in the effects of ArCCK1 and ArCCK2 on cardiac stomach preparations. To enable normalisation of the effects of the ArCCKP-derived peptides between experiments, the neuropeptide NGFFYamide was also tested on each preparation at a concentration of 100 nM (Figure 6a), and at this concentration the effects of ArCCK1 and ArCCK2 (1 µM) were not significantly different (Student t-test; P > 0.05) to the effect of NGFFYamide (data not shown).

**Figure 6.**
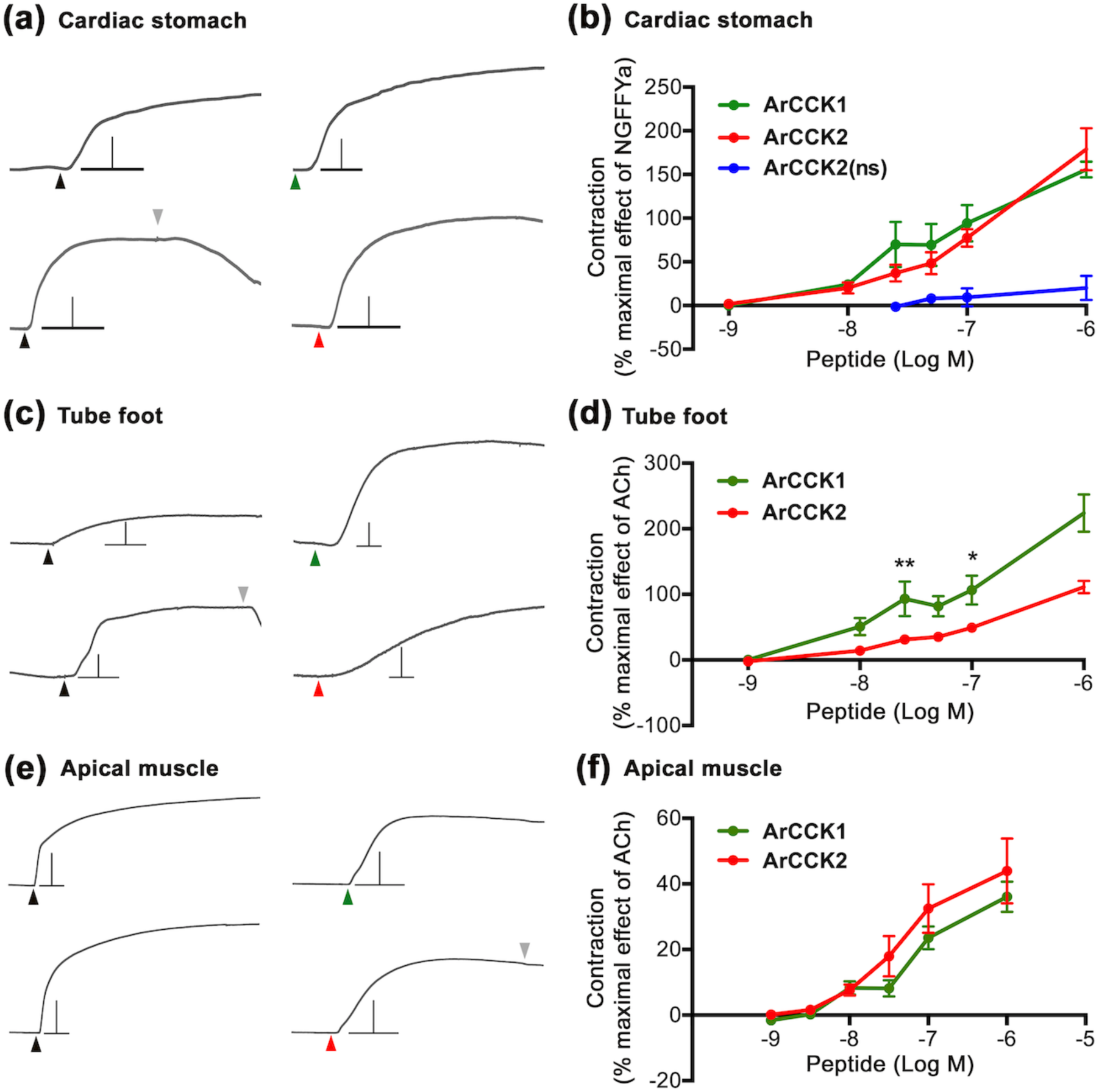
ArCCK1 and ArCCK2 cause concentration-dependent contraction of *in vitro* preparations of cardiac stomach, tube foot and apical muscle from *A. rubens*. **(a)** Representative recordings of the effects ArCCK1 (1 µM; green arrowhead) and ArCCK2 (1 µM; red arrowhead) in causing contraction of cardiac stomach preparations and compared with the effect of NGFFYamide (100 nM; black arrowhead) on the same preparation. The downward pointing grey arrowhead shows when a preparation was washed with seawater. Scale bar: vertical 0.5 mV; horizontal 1 min. **(b)** Concentration-response curves comparing the effects of ArCCK1, ArCCK2 and ArCCK2(ns) on cardiac stomach preparations. The effects of peptides (means ± s.e.m; n = 5 – 9) were normalized to the effect of 100 nM NGFFYamide (NGFFYa). **(c)** Representative recordings of the effects ArCCK1 (1 µM; green arrowhead) and ArCCK2 (1 µM; red arrowhead) in causing contraction of tube foot preparations and compared with the effect of acetylcholine (10 µM; black arrowhead) on the same preparation. The downward pointing grey arrowhead shows when a preparation was washed with seawater. Scale bar: vertical 0.08 mV; horizontal 1 min. **(d)** Concentration-response curves comparing the effects of ArCCK1 and ArCCK2 on tube foot preparations. The effects of peptides (means ± SEM; n = 8 - 10) were normalized to the effect of 10 µM acetylcholine (ACh). * indicates statistically significant differences between ArCCK1 and ArCCK2 when tested at concentrations of 25 nM and 100 nM (P < 0.01 and P < 0.05 respectively) as determined by 2-way ANOVA and Bonferroni’s multiple comparison test. **(e)** Representative recordings of the effects ArCCK1 (1 µM; green arrowhead) and ArCCK2 (1 µM; red arrowhead) in causing contraction of apical muscle preparations and compared to the effect of acetylcholine (10 µM; black arrowhead) on the same preparation. The downward pointing grey arrowhead shows when a preparation was washed with seawater. Scale bar: vertical 0.4 mV; horizontal 1 min. **(f)** Concentration-response curves comparing the effects of ArCCK1 and ArCCK2 on apical muscle preparations. The effects of peptides (means ± s.e.m; n = 20 - 23) were normalized to the effect of 10 µM acetylcholine (ACh).

Consistent with the effects of ArCCKP-derived neuropeptides on cardiac stomach preparations, ArCCK1 and ArCCK2 (1 nM to 1 µM) also caused concentration-dependent contraction of tube foot preparations (Figure 6c, d). Furthermore, comparison of the effects of ArCCK1 and ArCCK2 on tube feet using a 2-way ANOVA revealed significant differences, irrespective of concentration (P < 0.0001). In addition, Bonferroni’s multiple comparison test showed that ArCCK1 is significantly more effective than ArCCK2 when tested at concentrations of 25 nM and 100 nM (P < 0.01 and 0.05 respectively). Furthermore, ArCCK2(ns) peptide did not cause contraction of tube foot preparations *in vitro* (data not shown).

Although ArCCKP/ArCCK1 expression was not detected in the apical muscle (see above), we nevertheless tested ArCCK1 and ArCCK2 on this preparation because previous studies have revealed that other neuropeptides cause contraction (Tian et al. 2017) or relaxation (Cai et al. 2018, Lin et al. 2017a, Tinoco et al. 2018) of the apical muscle. Interestingly, consistent with the effects of ArCCKP-derived neuropeptides on cardiac stomach and tube foot preparations, both ArCCK1 and ArCCK2 caused contraction of apical muscle preparations (Figure 6e, f). However, by comparison with the effect of acetylcholine, which was tested at concentration of 10 µM to normalise effects of the peptides on different preparations, the contracting actions of ArCCK1 and ArCCK2 were only ∼40% of the effect of 10 µM ACh at the highest concentration tested (1 µM; Figure 6f). In contrast, the mean effects of ArCCK1 and ArCCK2 on tube foot preparations at 1 µM were ∼220% and ∼110% of the effect of 10 µM ACh (Figure 6d). No significant differences in the effects of ArCCK1 and ArCCK2 on apical muscle preparations were observed (2-way ANOVA, P > 0.05) and, as was observed with tube feet, ArCCK2(ns) had no effect on apical muscle preparations (data not shown).

### ArCCK1 and ArCCK2 trigger cardiac stomach retraction *in vivo*

Previous studies have revealed that the starfish neuropeptide NGFFYamide causes contraction of cardiac stomach preparations *in vitro* and triggers retraction of the everted cardiac stomach *in vivo* (Semmens et al. 2013). Because both ArCCK1 and ArCCK2 also cause contraction of cardiac stomach preparations *in vitro*, it was of interest to investigate if these neuropeptides also trigger retraction of the everted cardiac stomach *in vivo*. As reported previously (Semmens et al. 2013), cardiac stomach eversion was induced by immersing starfish in seawater containing 2% added MgCl_2_. In control experiments where starfish were injected with water, no retraction of the cardiac stomach was observed (Figure 7). However, injection of ArCCK1 or ArCCK2 (10 μl of 1 mM) triggered cardiac stomach retraction (P < 0.0001 for both peptides; 2-way ANOVA) (Figure 7a, b, d; **Videos 1, 2**), consistent with the contracting action of these peptides *in vitro*. ArCCK1 and ArCCK2 triggered cardiac stomach retraction in all animals tested but with some variability in the rate and extent of retraction. Consistent with the modest effect of ArCCK2(ns) on cardiac stomach preparations *in vitro* (Figure 6b), injection of ArCCK2(ns) (10 μl of 1 mM) did not trigger cardiac stomach retraction *in vivo* (Figure 7c).

**Figure 7.**
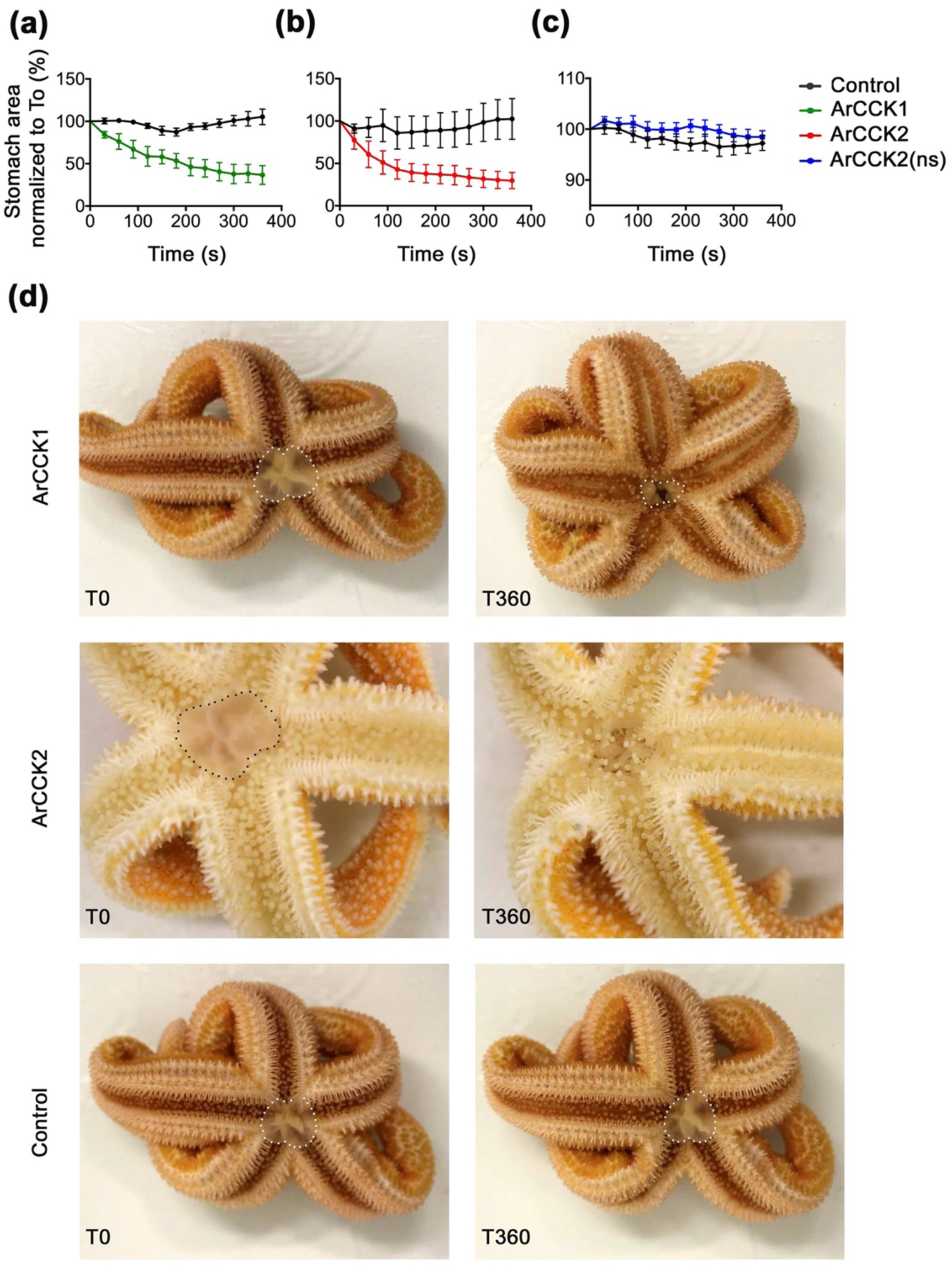
ArCCK1 and ArCCK2 trigger cardiac stomach retraction in *A. rubens*. The graphs compare experiments where starfish were first injected with vehicle (black line; 10 µl of distilled water) and then injected with **(a)** ArCCK1 (green line; 10 µl 1 mM), or **(b)** ArCCK2 (red line; 10 µl 1 mM), or **(c)** ArCCK2(ns) (blue line; 10 µl 1 mM). Stomach eversion was induced by placing starfish in seawater containing 2% MgCl_2_ and then the area of cardiac stomach everted (in 2D) at 30 s intervals (0-360 s) following injection of water (control) or peptide was measured, normalizing to the area of cardiac stomach everted at the time of injection (T0). Data (means ± s.e.m) were obtained from 6 (ArCCK1), 7 (ArCCK2) or 8 (ArCCK2(ns)) experiments. Both ArCCK1 and ArCCK2 cause retraction of the cardiac stomach, with >50% reduction in the area of cardiac stomach everted within 360 s (see videos 1 and 2), whereas ArCCK2(ns) has no effect. **(d)** Photographs from representative experiments showing that injection of ArCCK1 (10 µl 1 mM at T0) or ArCCK2 (10 µl 1 mM at T0) causes retraction of the everted cardiac stomach (marked with white or black dots), which is reflected in a reduction in the area everted after 360 s (T360). By way of comparison, in a control experiment injection with vehicle (10 µl of distilled water at T0) does not trigger cardiac stomach retraction.

### ArCCK1 and ArCCK2 inhibit feeding behaviour in *A. rubens*

The effects of ArCCK1 and ArCCK2 in triggering cardiac stomach retraction in *A. rubens* (see above) suggested that CCK-type signalling may have a physiological role in inhibition and/or termination of feeding behaviour in starfish, which would be consistent with the physiological roles of CCK-type neuropeptides in other taxa (Al-Alkawi et al. 2017, Downer et al. 2007, Kang et al. 2011, Maestro et al. 2001, Meyering-Vos and Muller 2007, Nachman et al. 1986b, Nässel et al. 2019, Rehfeld 2017, Roman et al. 2017, Wei et al. 2000, Yu et al. 2013a, Zels et al. 2015, Zhang et al. 2017). Therefore, we performed experiments to specifically investigate if ArCCK1 and ArCCK2 have inhibitory effects on starfish feeding behaviour on prey (mussels). Injection of ArCCK1 or ArCCK2 (10 µl of 1 mM) did not affect the time taken for starfish to make first contact with a mussel (time to touch; Figure 8a, b). However, compared to water-injected controls the mean time taken for starfish to adopt a feeding posture (time to enclose) was higher in neuropeptide-injected animals, reaching statistical significance (P < 0.05) with ArCCK2 treatment (P < 0.05; Figure 8d) but showing only a tendency to an increased time to enclose with ArCCK1 treatment (P = 0.0523; **Figure 9c**). This increased time to enclose was also reflected in an increased number of advances to touch the mussel in the starfish treated with ArCCK1 or ArCCK2 (data not shown). Moreover, by comparison with control starfish (water-injected), fewer starfish injected with ArCCK1 or ArCCK2 proceeded to initiation of a feeding posture after the first touch (P < 0.0001 and P < 0.01 for ArCCK1 and ArCCK2 respectively; Figure 8e, f). Another observation indicative of an inhibitory effect on feeding behaviour was that four and two of the starfish from ArCCK1- and ArCCK2-treated groups, respectively, did not initiate feeding on a mussel within the 5 h (300 min) observation period of the experiment, although feeding was commenced later and within 24 h of initiating the experiment.

**Figure 8.**
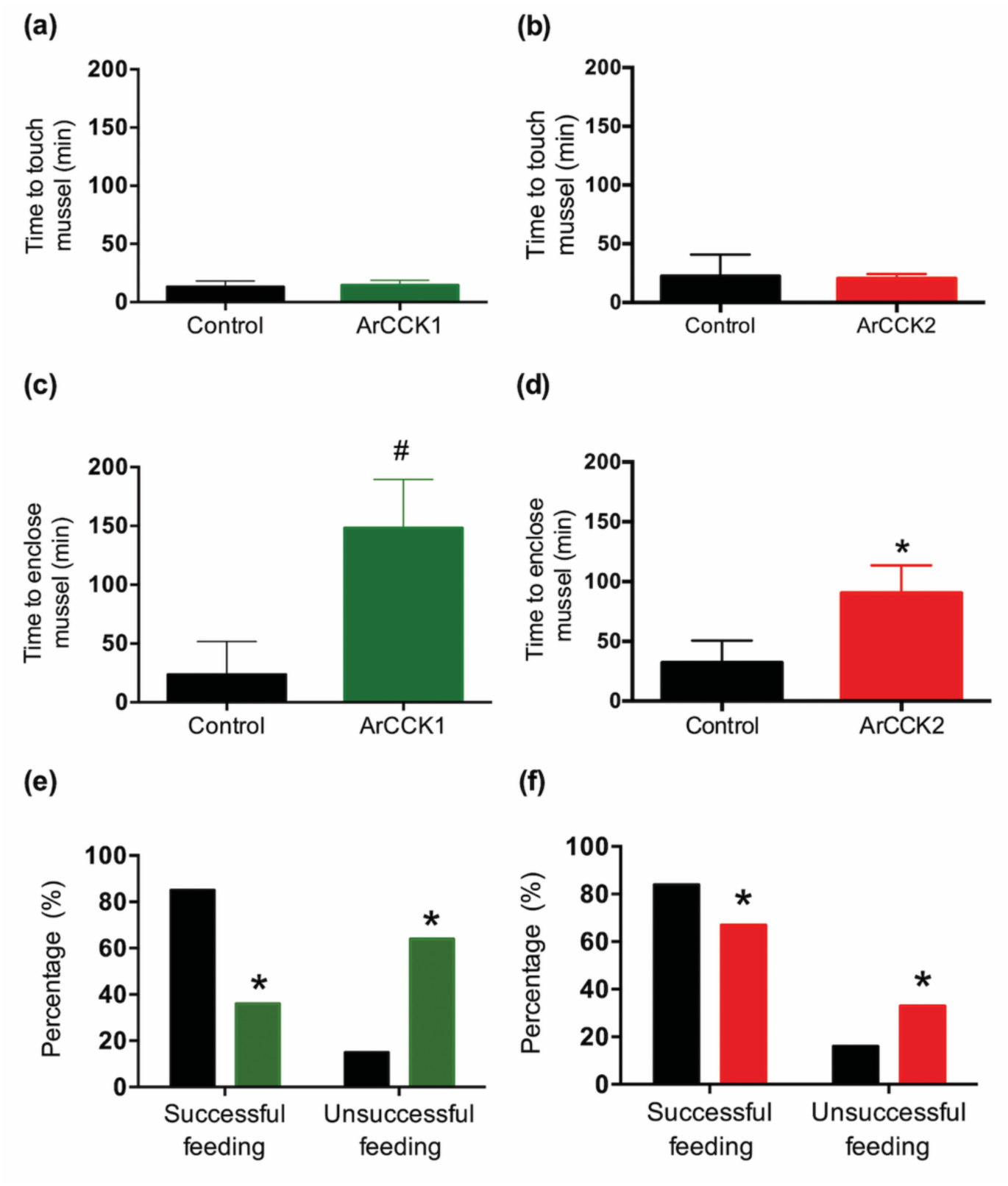
Effects of ArCCK1 and ArCCK2 on feeding behaviour in *A. rubens*. To investigate if injection of ArCCK1 and/or ArCCK2 affects feeding behaviour, starved animals were presented with a mussel as prey and then behaviour was observed. By comparison with vehicle-injected animals (control; 10 µl of distilled water; shown in black), injection of ArCCK1 **(a;** 10 µl of 1 mM; shown in green) or ArCCK2 (**b**; 10 µl of 1 mM; shown in red) had no effect on the time taken for starfish to make first contact with the mussel. However, by comparison with vehicle-injected animals (control; 10 µl of distilled water; shown in black) injection of ArCCK1 (**c**; 10 µl of 1 mM; shown in green) or ArCCK2 (**d**; 10 µl of 1 mM; shown in red) causes an increase in the time elapsed before starfish enclose a mussel. Data are expressed as means ± s.e.m (n = 13 for control- and 11 for ArCCK1-treated groups; n = 19 for control- and 19 for ArCCK2-treated groups). # indicates a nearly statistically significant difference (P = 0.0523) between vehicle-injected and ArCCK1-injected groups, as determined by two-tailed Mann-Whitney U-test. * indicates statistically significant differences (P < 0.05) between vehicle-injected and ArCCK2-injected groups, as determined by two-tailed Welch’s unequal variances t-test. Furthermore, injection of ArCCK1 (**e**; 10 µl of 1 mM; shown in green) or ArCCK2 (**f**; 10 µl of 1 mM; shown in red) causes a significant decrease in the percentage of starfish that initiate feeding after the mussel is touched for the first time, by comparison with vehicle-injected animals (control; 10 µl of distilled water; shown in black). * indicates statistically significant differences (P < 0.0001 and P < 0.01 for ArCCK1- and ArCCK2-treated groups respectively) between vehicle-injected and ArCCK1- or ArCCK2-injected groups, as determined by Fisher’s exact test.

## Discussion

The evolutionary origin of CCK/SK-type neuropeptide signalling has been traced to the common ancestor of the Bilateria, informed by the molecular characterisation of the orthologous CCK-type and SK-type neuropeptide signalling systems in chordates and protostomes, respectively (Bloom et al. 2019, Janssen et al. 2008, Johnsen and Rehfeld 1990, Kubiak et al. 2002, Mirabeau and Joly 2013, Monstein et al. 1993, Nachman et al. 1986a, Schwartz et al. 2018, Yu et al. 2013b, Yu and Smagghe 2014b). Here we report the first molecular and functional characterisation of a CCK/SK-type neuropeptide signalling system in a non-chordate deuterostome - the starfish *A. rubens* (phylum Echinodermata). This provides a key ‘missing link’ in our knowledge of the evolution and comparative physiology of CCK/SK-type signalling, complementing previously reported investigations of CCK-type signalling in chordates and SK-type signalling in protostome invertebrates (e.g. insects).

### Molecular characterisation of CCK-type signalling system in an echinoderm - the starfish *A. rubens*

Two CCK-type neuropeptides (ArCCK1 and ArCCK2) derived from the precursor protein ArCCKP were detected by mass spectrometry in *A. rubens* radial nerve cord extracts. An evolutionarily conserved feature of CCK-type neuropeptides is a tyrosine (Y) residue that is post-translationally modified by the addition of a sulphate group (Dufresne et al. 2006, Schwartz et al. 2018, Yu and Smagghe 2014a) and accordingly both ArCCK1 and ArCCK2 have a sulphated tyrosine. However, non-sulphated forms of these two peptides, ArCCK1(ns) and ArCCK2(ns), were also detected in *A. rubens* nerve cord extracts. Analysis of *A. rubens* neural transcriptome sequence data identified a GPCR (ArCCKR) that is an ortholog of CCK-type receptors that have been characterised in other taxa. Furthermore, heterologous expression of ArCCKR in CHO-K1 cells revealed that the sulphated forms of ArCCK1 and ArCCK2 are potent agonists for ArCCKR, whereas ArCCK2(ns) exhibited little or no agonist activity on this receptor. Therefore, we conclude that the sulphated neuropeptides ArCCK1 and ArCCK2 and the GPCR ArCCKR comprise the CCK-type neuropeptide signalling system in the starfish *A. rubens*. The requirement for the tyrosine residues in ArCCK1 and ArCCK2 to be sulphated in order that these peptides can act as potent ligands for ArCCKR is consistent with the properties of many CCK-type peptides in other taxa, including peptides that act as ligands for CCKR1 in mammals and SK-type peptides that act as ligands for the *Drosophila* receptor DSK-R1 (Dufresne et al. 2006, Kubiak et al. 2002). However, this is not a universal feature of CCK-type signalling; for example, non-sulphated CCK-type peptides act as potent ligands for CCK2R in mammals (Dufresne et al. 2006) and for CKR2a and CKR2b in the nematode *C. elegans* (Janssen et al. 2008).

Comparison of the sequences of ArCCK1 and ArCCK2 with the sequences of CCK-type peptides that have been identified in other taxa reveal a number of evolutionarily conserved features, including C-terminal amidation and the aforementioned tyrosine (Y) residue. The position of the tyrosine (Y) residue with respect to the C-terminal amide is variable amongst CCK-type peptides, with between five and seven intervening residues. Accordingly, in ArCCK1, ArCCK2 and CCK-type peptides from other echinoderms and hemichordates (collectively Ambulacraria) there are six intervening residues. In ArCCK1 the tyrosine (Y) residue is preceded by an aspartate (D) residue and in both ArCCK1 and ArCCK2 the tyrosine (Y) residue is followed by a glycine (G) residue; accordingly, a DYG motif is a feature of CCK-type peptides in many other taxa. ArCCK1, but not ArCCK2, has an N-terminal pyroglutamate in its mature form and this post-translational modification is also predicted or known to occur in some CCK-type peptides in other taxa – e.g. the SK1 and SK2 peptides from the cockroach *L. maderae* (Predel et al. 1999) and the CCK/SK-type peptides in the molluscs *Pinctata fucata* and *C. gigas* (Schwartz et al. 2018, Stewart et al. 2014). The C-terminal residue of ArCCK2 is a phenylalanine residue (F) and in this respect ArCCK2 is like the majority of CCK-type peptides that have been identified in other taxa. It is noteworthy, therefore, that the C-terminal residue of ArCCK1 is a tryptophan (W) residue, which is also a feature of an ArCCK1-like peptide sequence in another starfish species – the crown-of-thorns starfish *Acanthaster planci* (Smith et al. 2017). Furthermore, this unusual feature of one of the two CCK-type peptides that occur in starfish species appears to be unique to this class of echinoderms (the Asteroidea) because CCK-type peptides that have been identified in other echinoderms all have the more typical C-terminal phenylalanine (F) residue (Chen et al. 2019, Zandawala et al. 2017). Interestingly, however, it is not completely unique to starfish because in the bivalve mollusc *C. gigas* there are two CCK-type peptides, one of which has a C-terminal phenylalanine (F) and another that has a tryptophan (W) residue (Schwartz et al. 2018). Thus, it appears that CCK-type neuropeptides with a C-terminal tryptophan (W) residue may have evolved independently in starfish and in the bivalve mollusc *C. gigas*.

### Functional characterization of CCK-type neuropeptides in an echinoderm - the starfish *A. rubens*

Having identified the molecular components of a CCK-type neuropeptide signalling system in *A. rubens*, comprising the sulphated neuropeptides ArCCK1 and ArCCK2 and their receptor ArCCKR, our next objective was to gain insights into the physiological roles of CCK-type neuropeptides in starfish. Using mRNA *in situ* hybridisation and immunohistochemistry, we examined the anatomical expression patterns of ArCCKP transcripts and one of the neuropeptides derived from ArCCKP (ArCCK1), respectively. This revealed a widespread pattern of expression, including the central nervous system (CNS), tube feet and other body wall associated structures and the digestive system, as discussed below. This pattern of expression can be interpreted with reference to the anatomy of the starfish nervous system (Cobb 1970, Mashanov et al. 2016, Smith 1937), digestive system (Anderson 1953, 1954) and body wall (Blowes et al. 2017) and in comparison with the distribution of other neuropeptides in *A. rubens* (Cai et al. 2018, Elphick et al. 1995, Lin et al. 2017a, Lin et al. 2018, Moore and Thorndyke 1993, Odekunle et al. 2019, Tian et al. 2017, Tinoco et al. 2018, Yáñez-Guerra et al. 2018, Zhang et al. 2020).

The starfish CNS consists of radial nerve cords and a circumoral nerve ring, where cells expressing ArCCKP/ArCCK1 were revealed in both the ectoneural and hyponeural regions in common with other neuropeptides such as the calcitonin-type neuropeptide ArCT (Cai et al. 2018), pedal peptide/orcokinin-type peptides (Lin et al. 2017a, Lin et al. 2018); the gonadotropin-releasing hormone-type peptide ArGnRH (Tian et al. 2017); and the somatostatin-type peptide ArSS2 (Zhang et al. 2020). The presence of an extensive network of immunostained fibres in the neuropile of the ectoneural region is consistent with neuronal expression of ArCCKP/ArCCK1. Furthermore, regional variation in the density of ArCCK1-ir fibres in the ectoneural neuropile was observed. The ectoneural region is thought to contain sensory neurons, interneurons and motor neurons (Brusca 2017, Cobb 1970, Smith 1937) but the functional identity of neuronal cell types and the neural circuitry are not known. Therefore, it is not possible at present to determine the functional properties of ArCCK-ir neurons in the ectoneural region of the CNS. However, the hyponeural region of the starfish CNS only comprises motoneurons and the projection pathways of the axons of these motoneurons have been reported (Lin et al. 2017a, Smith 1950). Thus, the axons of hyponeural motor neurons project around the tube feet and coalesce as a fibre bundle known as the lateral motor nerve (Lin et al. 2017a, Smith 1937). Consistent, with the expression of ArCCKP in hyponeural cells, ArCCK1-ir fibres can be seen in the lateral motor nerves, as has been previously reported for other *A. rubens* neuropeptides (Cai et al. 2018, Lin et al. 2017a, Lin et al. 2018, Zhang et al. 2020). Branches of the lateral motor nerves project into the coelomic lining of the body wall in starfish (Smith 1937, 1950) and accordingly ArCCK1-ir fibres were observed in coelomic lining of the body wall in *A. rubens*. Furthermore, the presence of ArCCK1-ir fibres in inter-ossicular tissue of the body wall in *A. rubens* suggests that ArCCKP-expressing hyponeural cells may include motoneurons that innervate inter-ossicular muscles and/or inter-ossicular mutable collagenous tissue (Blowes et al. 2017).

ArCCKP/ArCCK1 is widely expressed in the digestive system of *A. rubens*, including the oesophagus, peristomial membrane, cardiac stomach, pyloric stomach, pyloric duct, pyloric caeca, intestine and rectal caeca. Consistent with the expression of ArCCKP/ArCCK1 in the cardiac stomach, ArCCK1 and ArCCK2 cause concentration-dependent contraction of cardiac stomach preparations *in vitro*. ArCCKP/ArCCK1-expressing cells are present in the mucosal wall of the cardiac stomach and a dense network of ArCCK1-ir fibres is present in the basiepithelial nerve plexus of the cardiac stomach, particularly in the highly folded lateral pouches of the cardiac stomach. Therefore, CCK-type neuropeptides released by fibres in the basiepithelial nerve plexus of the cardiac stomach may diffuse across an intervening thin layer of collagenous tissue to cause contraction of visceral muscle. Other peptides that cause cardiac stomach contraction *in vitro* have been identified in *A. rubens*, which include NGFFYamide (Semmens et al. 2013), the GnRH-type peptide ArGnRH and the corazonin-type neuropeptide ArCRZ (Tian et al. 2017). In comparison with effects of NGFFYamide on cardiac stomach preparations *in vitro* (Semmens et al. 2013), ArCCK1 and ArCCK2 exhibit lower potency but higher efficacy. In comparison with effects of ArGnRH and ArCRZ on cardiac stomach preparations *in vitro* (Tian et al. 2017), ArCCK1 and ArCCK2 exhibit higher potency and efficacy. These observations on the *in vitro* effects ArCCK1 and ArCCK2 on cardiac stomach preparations provided a basis for examining the *in vivo* effects of these peptides on feeding related processes in *A. rubens*.

Starfish exhibit one of the most remarkable feeding behaviours in the animal kingdom – they evert their stomach out of their mouth and digest large prey externally, and once the prey has been digested, the stomach is withdrawn. Extra-oral feeding in starfish requires relaxation of muscle in the wall of the cardiac stomach and in intrinsic and extrinsic retractor strands to enable stomach eversion. Then when external digestion and ingestion of prey tissue is completed, contraction of the musculature enables cardiac stomach retraction (Anderson 1954). Previous studies have identified neuropeptides that cause relaxation of the cardiac stomach *in vitro* and trigger cardiac stomach eversion when injected *in vivo*; for example, the SALMFamide neuropeptide S2 (Melarange et al. 1999), the vasopressin/oxytocin-type neuropeptide asterotocin (Odekunle et al. 2019) and the somatostatin-type neuropeptide ArSS2 (Zhang et al. 2020). Conversely, NGFFYamide has been identified as a neuropeptide in *A. rubens* that triggers contraction of the cardiac stomach *in vitro* and retraction of the everted stomach when injected *in vivo* (Semmens et al. 2013). The *in vitro* effects of ArCCK1 and ArCCK2 in causing contraction of the cardiac stomach and the presence of ArCCK1-ir fibres in the basiepithelial nerve plexus of the cardiac stomach and in retractor strands suggest that CCK-type peptides may participate in mechanisms of cardiac stomach retraction physiologically. Accordingly, injection of 10 µl of 1 mM ArCCK1 or 1 mM ArCCK2 triggered partial or complete retraction of the cardiac stomach within a test period of six minutes. These experiments indicate that CCK-type peptides may be involved in physiological mechanisms of cardiac stomach retraction in starfish. Furthermore, to investigate the importance of tyrosine (Y) sulphation for the bioactivity of CCK-type peptides in *A. rubens*, we also tested the non-sulphated peptide ArCCK2(ns) on cardiac stomach preparations. By comparison with ArCCK1 and ArCCK2, ArCCK2(ns) had a very modest contracting effect on cardiac stomach preparations *in vitro* and did not trigger cardiac stomach retraction *in vivo*. This is consistent with low potency of ArCCK2(ns) as an agonist on ArCCKR expressed in CHO cells and indicative that non-sulphated CCK-type peptides are not bioactive physiologically in starfish.

The *in vivo* effect of CCK-type neuropeptides in triggering cardiac stomach retraction is indicative of a role in physiological mechanisms that control termination of feeding behaviour in starfish. By way of comparison, the neuropeptide NGFFYamide that triggers retraction of the everted stomach of *A. rubens* also causes a significant delay in the onset of feeding on prey (mussels) when injected *in vivo* (Tinoco et al. 2018). Accordingly, here we observed that starfish injected with CCK-type neuropeptides took longer to enclose prey compared to control animals injected with water. Furthermore, in animals injected with CCK-type neuropeptides the proportion of starfish that successfully consumed prey was fewer than in control animals that were injected with water. Thus, collectively these findings indicate that CCK-type signalling acts as a physiological regulator that inhibits and/or terminates feeding behaviour in starfish.

Investigation of the *in vitro* pharmacological effects of neuropeptides in starfish has revealed that some peptides that act as contractants or relaxants of the cardiac stomach also cause contraction or relaxation, respectively, of two other muscular tissues/organs – tube feet and the body wall associated apical muscle. For example, the GnRH-type peptide ArGnRH and the corazonin-type peptide ArCRZ cause contraction of all three preparations (Tian et al. 2017) and the SALMFamide-type neuropeptides S1 and S2 and the pedal peptide/orcokinin-type peptide ArPPLN1b (starfish myorelaxant peptide) cause relaxation of all three preparations (Lin et al. 2017a, Melarange and Elphick 2003).

Informed by detection of ArCCKP/ArCCK1 expression in the tube feet of *A. rubens*, we tested ArCCK1 and ArCCK2 on *in vitro* preparations of these locomotory organs and found that both peptides cause dose-dependent contraction. ArCCKP-expressing cells are present at the base of tube foot podium proximal to the junction with the radial nerve cord or marginal nerve and also in the disk region proximal to the basal nerve ring. Consistent with this pattern of precursor protein expression, ArCCK1-ir was revealed in the basiepithelial nerve plexus and in the basal nerve ring. The basiepithelial nerve plexus is separated from the tube foot muscle layer by a thin layer of collagenous tissue and therefore, consistent with mechanisms proposed for control tube foot myoactivity by the neurotransmitter acetylcholine (Florey and Cahill 1980, Florey et al. 1975), CCK-type peptides released by nerve processes in the basiepithelial nerve plexus may diffuse across the collagenous tissue layer to cause contraction of the longitudinally orientated tube foot muscle *in vivo*. Furthermore, from a behavioural perspective, CCK-type neuropeptides may participate in neural mechanisms controlling tube foot retraction during locomotor activity and/or for generation of force when the collective pulling power of tube feet is used by starfish to prise apart the valves of prey such as mussels (Lavoie 1956).

Although expression of ArCCKP/ArCCK1 was not detected in the apical muscle, ArCCK1-ir fibres were revealed proximally in the coelomic lining of the body wall. Therefore, we also tested ArCCK1 and ArCCK2 on *in vitro* preparations of the apical muscle from *A. rubens* and found that both peptides caused dose-dependent contraction. However, by comparison with their effects on tube foot preparations, the effects of ArCCK1 and ArCCK2 on apical muscle preparations were quite modest. Thus, the effects of 1 µM ArCCK1 or ArCCK2 on apical muscle preparations were ∼40% of the effect of 10 µM ACh, whereas the effects of 1 µM ArCCK1 or ArCCK2 on tube foot preparations were, respectively, 220% and 110% of the effect of 10 µM ACh. Finally, it is noteworthy that ArCCK1-ir fibres were revealed in the inter-ossicular tissue that contains muscles and mutable collagenous tissue that interlink the calcite ossicles of the body wall endoskeleton (Blowes et al. 2017). Therefore, CCK-types neuropeptides in *A. rubens* may also have physiological roles in regulating the contractile state of inter-ossicular muscles and/or the stiffness of inter-ossicular mutable collagenous tissue.

### Comparative and evolutionary physiology of CCK/SK-type neuropeptide signalling in the Bilateria

Our *in vitro* pharmacological analysis of the effects of ArCCK-type neuropeptides in *A. rubens* revealed a myoexcitatory action, consistent with findings from previous studies on vertebrates and protostome invertebrates. For example, CCK causes pyloric and gallbladder contraction in mammals (Gutiérrez et al. 1974, Rehfeld 2017, Shaw and Jones 1978, Vizi et al. 1973) and gut contraction in other vertebrates (Tinoco et al. 2015). Accordingly, SK-type peptides also have myoexcitatory effects in insects (Al-Alkawi et al. 2017, Maestro et al. 2001, Marciniak et al. 2011, Nachman et al. 1986b, Nichols 2007, Palmer et al. 2007, Predel et al. 2001). However, CCK-type peptides are not exclusively myoexcitatory because, for example, CCK causes relaxation of the proximal stomach in mammals; however, this myoinhibitory effect of CCK is indirect and mediated by vagal and splanchnic afferents (Takahashi and Owyang 1999). Accordingly, both inhibitory and excitatory effects of CCK-8 on the stomach of a non-mammalian vertebrate, the rainbow trout *Oncorhynchus mykiss*, have been reported (Olsson et al. 1999) and a CCK/SK-type peptide causes a decrease in the frequency of hindgut contraction in the mollusc *C. gigas* (Schwartz et al. 2018). Furthermore, and directly relevant to this study, it is has been reported that mammalian CCK-8 causes *in vitro* relaxation of intestine preparations from the sea cucumber of *Holothuria glaberrima* (García-Arrarás et al. 1991). With the determination of the amino-acid sequences of native CCK-type neuropeptides in sea cucumbers (Chen et al. 2019, Zandawala et al. 2017), it will now be possible to specifically investigate their pharmacological effects in these animals to make direct comparisons with the findings reported here for starfish.

As discussed below, CCK/SK-type neuropeptides are perhaps best known for their roles as inhibitory regulators of feeding. However, in common with other neuropeptides, they are pleiotropic in their physiological roles. Thus, linked to regulation feeding, CCK/SK-type neuropeptides stimulate secretion of gastric acid and/or digestive enzymes in mammals, insects, nematodes, ascidians and molluscs (Bevis and Thorndyke 1981, Chen et al. 2004, Harper and Raper 1943, Harshini et al. 2002b, Janssen et al. 2008, Nachman et al. 1997, Shaw and Jones 1978, Thorndyke and Bevis 1984, Zels et al. 2015). Accordingly, expression of CCK-type peptides by cells in several regions of the digestive system in *A. rubens* may be indicative of a similar role in starfish. Furthermore, CCK precursor transcripts are detected in rat spinal motoneurons (Cortés et al. 1990) and SK-type neuropeptides act as positive growth regulators for neuromuscular junction formation and promote locomotion in larval *Drosophila* (Chen and Ganetzky 2012). In this context, it is noteworthy that CCK-type neuropeptides cause contraction of body wall associated muscles (apical muscle) and organs (tube feet) in starfish. Accordingly, the expression of CCK-type peptides in hyponeural motoneurons, the lateral motor nerve and inter-ossicular tissue of the body wall of *A. rubens* may reflect evolutionarily ancient and conserved roles of CCK-type neuropeptides as regulators of skeletal muscle function. It is also noteworthy that CCK is one of the most abundantly expressed neuropeptides in the cortex of the mammalian brain, where it is expressed by sub-populations of GABAergic interneurons and acts as a multi-functional molecular switch to regulate the output of cortical neuronal circuits (Lee and Soltesz 2011). Furthermore, evidence that expression of CCK in GABAergic neurons is an evolutionarily ancient association was provided by a recent study reporting co-localisation of CCK-8 and GABA in several different neuronal populations in the brain of the sea lamprey *Petromyzon marinus* (Sobrido-Cameán et al. 2020). GABA-immunoreactive neurons have been revealed in the ectoneural region of the radial nerve cord in *A. rubens* (Newman and Thorndyke 1994) and sub-populations of these neurons may correspond with cells expressing CCK-type neuropeptides reported in this study. However, the inaccessibility and small size of these neurons may preclude investigation of their electrophysiological properties.

The key functional insights from this study are our observations that in the starfish *A. rubens* CCK-type neuropeptides trigger cardiac stomach contraction and retraction and induce a delay in the onset of feeding and a reduction in predation. These findings are of general interest because of the previously reported evidence that CCK/SK-type neuropeptides mediate physiological mechanisms of satiety and/or regulate feeding behaviour in vertebrates, insects and the mollusc *C. gigas* (Al-Alkawi et al. 2017, Downer et al. 2007, Himick and Peter 1994, Kang et al. 2011, Maestro et al. 2001, Meyering-Vos and Muller 2007, Nachman et al. 1986a, Nachman et al. 1986b, Nässel and Zandawala 2019, Rehfeld 2017, Roman et al. 2017, Schwartz et al. 2018, Wei et al. 2000, Yu et al. 2013a, Yu and Smagghe 2014b, Zels et al. 2015, Zhang et al. 2017). Furthermore, insights into the mechanisms by which CCK/SK-type neuropeptides regulate feeding behaviour in mammals and insects have been obtained. In mammals CCK released by intestinal endocrine cells acts on vagal afferents, which is thought to then lead to activation of calcitonin-gene related peptide (CGRP)-expressing neurons in the parabrachial nucleus that supress feeding and inhibition of Agouti-related peptide (AgRP)-expressing hypothalamic neurons that promote feeding (Beutler et al. 2017, Essner et al. 2017). In *Drosophila*, SK-type neuropeptides are expressed by a sub-population median neurosecretory cells in the brain that also produce insulin-like peptides and results from a variety of experimental studies indicate that release of SK-type neuropeptides by these neurons induces satiety in both larval and adult flies (Nässel and Williams 2014). Thus, an evolutionarily conserved physiological role of CCK/SK-type neuropeptides as inhibitory regulators of feeding are mediated by different mechanisms in mammals and insects, which may reflect evolutionary divergence in the anatomy of these taxa. It is interesting, therefore, that in the unique context of the evolutionary and developmental replacement of bilateral symmetry with pentaradial symmetry in adult echinoderms, an ancient role of CCK/SK-type neuropeptides as inhibitory regulators of feeding-related processes has been retained in starfish. Furthermore, because feeding in starfish is accomplished by stomach eversion, there is a direct link between the action of CCK-type neuropeptides on the gastro-intestinal system and inhibition/termination of feeding behaviour. Thus, our findings from starfish reported here for CCK/SK-type neuropeptides and previously for other neuropeptides (Cai et al. 2018, Odekunle et al. 2019, Tian et al. 2017, Tinoco et al. 2018, Zhang et al. 2020) reveal how ancient roles of neuropeptide signalling systems have been preserved in spite of unique and radical evolutionary and developmental changes in the anatomy of echinoderms amongst bilaterian animals.

## Material and methods

### Animals

Adult starfish (*Asterias rubens*, Linnaeus, 1758) were collected at low tide near Margate (Kent, UK) or obtained from a fisherman based at Whitstable (Kent, UK). The starfish were maintained in a circulating seawater aquarium under a 12 h −12 h of light - dark cycle (lights on at 8 a.m.) at a temperature of ∼12°C and salinity of 32 ‰, located in the School of Biological & Chemical Sciences at Queen Mary University of London. Animals were fed on mussels (*Mytilus edulis*) that were collected at low tide near Margate (Kent, UK). Additionally, juvenile specimens of *A. rubens* (diameter 0.5 - 1.5 cm) used for anatomical studies were collected at the University of Gothenburg Sven Lovén Centre for Marine Infrastructure (Kristineberg, Sweden).

### Cloning of a cDNA encoding the *Asterias rubens* CCK-type precursor ArCCKP

A cDNA encoding the ArCCK precursor (ArCCKP), including 5’ and 3’ untranslated regions (UTR) and the complete open reading frame (ORF), was amplified by PCR (Phusion High-Fidelity PCR Master Mix, NEB, Hitchin, Hertfordshire, UK) using specific oligonucleotide primers (5’-TCGCTACTGTTTCTCTCGCA-3’ and 5’-AAAGGCGTCAACAACTGCTT-3’), which were designed using Primer3 software (http://bioinfo.ut.ee/primer3-0.4.0/) with reference to the ArCCKP transcript sequence (contig 1124413; GenBank accession number KT601716) obtained from *A. rubens* radial nerve cord transcriptome data (Semmens et al. 2016). The PCR product was gel-extracted and purified (QIAquick Gel Extraction Kit, Qiagen, Manchester, UK) before being blunt-end cloned into pBluescript SKII (C) (Agilent Technologies, Stockport, Cheshire, UK) or Zero Blunt® Topo PCR (ThermoFisher Scientific; Waltham, MA, USA) vectors. The clones were sequenced (Eurofins Genomics GmbH, Ebersberg, Germany) using the T7 and T3 sequencing primer sites.

### Localisation of ArCCKP expression in *A. rubens* using mRNA *in situ* hybridization

To enable visualisation of ArCCKP transcripts in *A. rubens* using mRNA *in situ* hybridization, digoxygenin-labelled RNA probes were synthesised. Zero Blunt® Topo or pBluescript SKII (+) vectors containing the ArCCKP cDNA were purified (Qiagen Maxiprep, Qiagen, Manchester, UK) and 5 μg of the vector was linearized using restriction enzymes (NEB, Hitchin, Hertfordshire, UK). Linearized vector containing the ArCCKP cDNA were cleaned using phenol-chloroform (Sigma-Aldrich, Gillingham, UK) and chloroform-isomylalcohol (Sigma-Aldrich, Gillingham, UK) extractions and then precipitated using 1/10 volume of 3 M sodium acetate and 2.5 volumes of 100% ethanol (Honeywell^TM^, Fisher Scientific UK Ltd, Loughborough, UK) at −80°C. The pellet was washed with 70% ice-cold ethanol before air drying and re-suspending in autoclaved water. Sense and antisense RNA probes were synthesized using digoxigenin nucleotide triphosphate (DIG-NTP) mix (Roche, Mannheim, Germany), 5x transcription buffer (NEB, Hitchin, Hertfordshire, UK), 0.2 M dithiothreitol (DTT) (Promega, Madison, USA), placental ribonuclease inhibitor (10 U/μl) (Promega, Madison, USA) and T7 polymerase (50 U/μl) or T3 polymerase (50 U/μl) (NEB, Hitchin, Hertfordshire, UK) with 1 μg of linearised vector containing the ArCCKP cDNA. Template DNA was then digested with RNase free DNase (NEB, Hitchin, Hertfordshire, UK). RNA probes were stored in 25% formamide/2x saline-sodium citrate (25% FA/2x SSC; VWR Chemicals, Leicestershire, UK) at −20°C for long-term storage.

To prepare specimens of *A. rubens* for mRNA *in situ* hybridisation, animals were fixed by immersion in 4% paraformaldehyde (PFA; Sigma-Aldrich, Gillingham, UK) in phosphate-buffered saline (PBS) overnight at 4°C. Specimens were washed in PBS, dissected and placed in Morse’s solution (10% sodium citrate; 20% formic acid in autoclaved water) to enable decalcification of ossicles in the body wall of starfish. Decalcified specimens were then washed in autoclaved water, dehydrated through a graded ethanol series and then immersed in xylene (Honeywell, Fisher Scientific UK Ltd, Loughborough, UK) before being embedded in paraffin wax. 14 μm sections of *A. rubens* arms and central disk were prepared using a RM 2145 microtome (Leica Microsystems [UK], Milton Keynes, UK). Sections were collected on poly-L-lysine coated slides (VWR Chemicals, Lutterworth, Leicestershire, UK) that had been placed on a hot plate and covered with autoclaved water. Slides were left to dry before proceeding with probe hybridization and immunodetection.

Slides were kept at 60°C for 1 hour to allow excess wax to melt before leaving to cool at room temperature. Sections were then deparaffinised in xylene and hydrated through a graded ethanol series before being washed in PBS. Sections were then post-fixed in 4% PFA/PBS before washing with buffer containing Proteinase K (PK; Qiagen UK Ltd, Manchester, UK) (1 µg/ml PK, 50 mM Tris-HCl [pH 7.5]; 6.25 mM EDTA in autoclaved water; Thermo Fisher Scientific, Oxford, UK) at 37°C for 12 minutes. Sections were then post-fixed in 4% PFA/PBS before washing with PBS/Tween 0.1% and then acetylated (1.325% triethanolamine [pH 7-8]; 0.25% acetic anhydride; 0.175% HCl in autoclaved water; VWR Chemicals, Lutterworth, UK) for 10 minutes. Sections were washed in PBS/0.1% Tween-20 and in 5x SSC. Then sections were dried, placed in a humidified chamber and covered with hybridisation buffer (50% formamide; 5x SSC; 500 μg/ml yeast total RNA; 50 μg/ml heparin; 0.1% Tween-20 in autoclaved water) at room temperature for 2 hours. ArCCK precursor sense and anti-sense probes (500-1000 ng/ml) were denatured in hybridisation buffer at 80°C and placed on ice before adding remaining hybridisation buffer and applying 100 μl probe solution per slide. Slides were covered with a piece of Parafilm (Bemis, Terre Haute, IN, USA) and then placed in a humidified chamber at 65°C overnight. Sections were then washed in 0.2x SSC at 65°C, in 0.2x SSC at room temperature and equilibrated in buffer B1 (10 mM Tris-HCl [pH 7.5]; 150 mM NaCl in autoclaved water). Sections were covered in buffer B1/5% goat serum and placed in a humidified chamber at room temperature for 2 hours. Sections were then dried and covered in an alkaline phosphatase (AP)-conjugated anti-DIG antibody (1:3000; Roche, Mannheim, Germany) in buffer B1/2.5% goat serum at 4°C overnight. Slides were washed in buffer B1 and then equilibrated in buffer B3 (100 mM Tris-HCl [pH 9.5]; 100 mM NaCl; 50 mM MgCl2 in autoclaved water). Sections were then covered in buffer B3/0.1% Tween-20 with nitro-blue tetrazolium chloride (NBT; Sigma-Aldrich, Gillingham, UK) (75 mg/ml in 70% dimethylformamide) and 5-bromo-4-chloro-3’-indolyphosphate p-toluidine salt (BCIP; Sigma-Aldrich, Gillingham, UK) substrate solution (50 mg/ml BCIP in autoclaved water) until strong staining was observed. The slides were washed in distilled water to stop the staining reaction and then were dried on a hot plate before rinsing in 100% ethanol and Histo-Clear (National Diagnostics, Fisher Scientific UK Ltd, Loughborough, UK). Sections were mounted with a coverslip on HistoMount solution (National Diagnostics, Fisher Scientific UK Ltd, Loughborough, UK) for long-term storage.

### Mass spectrometry

Extracts of *A. rubens* radial nerve cords were prepared and analysed using mass spectrometry (NanoLC-ESI-MS/MS), as described in detail previously for the *A. rubens* relaxin-like gonad stimulating peptide, which contains disulphide bridges (Lin et al. 2017b). Aliquots of radial nerve cord extract were not treated with trypsin but were subjected to reduction to break disulphide bridges (using 100 mM dithiothreitol; Sigma Aldrich, Gillingham, UK) followed by alkylation of cysteine residues (using 200 mM iodoacetamide; Sigma Aldrich. Gillingham, UK). Raw data were converted to Mascot generic format using MSConvert in ProteoWizard Toolkit (v. 3.0.5759) (Kessner et al. 2008). MS spectra were searched with Mascot engine (Matrix Science, v. 2.4.1) (Nesvizhskii et al. 2003) against a database comprising 40 *A. rubens* neuropeptide precursor proteins, including ArCCKP (Semmens et al. 2016), all proteins in GenBank from species belonging to the family Asteriidae and the common Repository of Adventitious Proteins Database (http://www.thegpm.org/cRAP/index.html). A no-enzyme search was performed with up to two missed cleavages and carbamidomethyl as a fixed modification. Post-translational amidation of C-terminal glycine residues, pyroglutamylation of N-terminal glutamine residues, sulphation of tyrosine residues and oxidation were included as variable modifications. Precursor mass tolerance was 10 ppm and product ions were searched at 0.8 Da tolerances.

Scaffold (version Scaffold_4.6.1, Proteome Software Inc.) was used to validate MS/MS based peptide and protein identifications. Peptide identifications were accepted if they could be established at greater than 95.0% probability by the Scaffold Local FDR algorithm. Protein identifications were accepted if they could be established at greater than 95.0% probability and contained at least two identified peptides.

### Alignment of the *A. rubens* CCK-type neuropeptides ArCCK1 and ArCCK2 with CCK-type peptides from other taxa

Having used mass spectrometry to confirm the structures of the mature peptides ArCCK1 and ArCCK2 that are derived from ArCCKP, the sequences of ArCCK1 and ArCCK2 were aligned with CCK-type peptides from other taxa to investigate the occurrence of evolutionarily conserved residues. The alignment was generated using MAFFT (version 7) with the following parameters (BLOSUM62, 200 PAM/K=2) and highlighted using BOXSHADE (http://www.ch.embnet.org/software/BOX_form.html) using 70% conservation as a minimum for highlighting. The accession numbers for the sequences used for this analysis are shown in Figure 1 – source data 1.

### Identification a transcript encoding an *A. rubens* CCK-type receptor

A transcript encoding an *A. rubens* CCK-type receptor (ArCCKR) was identified by tBLASTn analysis of the *A. rubens* radial nerve cord transcriptome data (Semmens et al. 2016), using SequenceServer [https://www.sequenceserver.com; (Priyam et al. 2015)] and a *Strongylocentrotus purpuratus* CCK-type receptor (Accession number XP_782630.3) as the query sequence. To investigate in more detail the relationship of ArCCKR with CCK-type receptors that have been identified in other taxa, phylogenetic analyses were performed using the maximum-likelihood method. The sequences of ArCCKR and CCK-type receptors from a variety of taxa were aligned using MUSCLE (iterative, 10 iterations, UPGMB as clustering method) (Edgar 2004). The maximum-likelihood tree was generated using IQ-tree web server [1000 bootstrap replicates, LG+F+I+G4 substitution model; (Trifinopoulos et al. 2016)]. The accession numbers of the protein sequences that were used for this analysis are listed in Figure 2 – source data 1.

### Pharmacological characterization of ArCCKR

To enable testing of ArCCK1 and ArCCK2 as candidate ligands for ArCCKR, a full-length cDNA encoding ArCCKR was synthesized by GenScript (Piscataway, NJ, USA) and cloned into pcDNA 3.1+ vector (Invitrogen^TM^, ThermoFisher Scientific; Waltham, MA, USA). A partial Kozak translation initiation sequence (CACC) was introduced upstream to the start codon (ATG). Chinese hamster ovary (CHO)-K1 cells stably expressing the mitochondrial targeted calcium-sensitive bioluminescent reporter GFP-aequorin fusion protein (G5A) were used as a heterologous expression system for ArCCKR. Cells were cultured, co-transfected with the pcDNA 3.1+ vector containing the ArCCKR cDNA sequence and plasmids encoding the promiscuous human G-protein G_α_16. Then bioluminescence-based receptor assays were performed, as described previously for *A. rubens* luqin-type receptors (Yáñez-Guerra et al. 2018).

CCK-type peptides in other taxa have a sulphated tyrosine residue that is important for bioactivity and therefore the ArCCK1 and ArCCK2 peptides were synthesized (Peptide Protein Research Ltd, Fareham, UK) with sulphated tyrosine residues: pQSKVDDY(SO_3_H)GHGLFW-NH_2_ (ArCCK1), and GGDDQY(SO_3_H)GFGLFF-NH_2_ (ArCCK2). Furthermore, to assess the requirement of tyrosine sulphation for receptor activation and bioactivity, a non-sulphated form of ArCCK2 was also synthesized: GGDDQYGFGLFF-NH_2_ [ArCCK2(ns)]. The peptides were diluted in distilled water and tested as candidate ligands for ArCCKR at concentrations ranging from 3 x 10^−17^ M to 10^−4^ M. Concentration-response data were determined as a percentage of the highest response for each peptide (100% activation). EC_50_ values were calculated from concentration-response curves based on 4 to 6 independent transfections and averaging 2 - 3 replicates in each transfection using Prism 6.0 (GraphPad software, La Jolla, CA). Cells transfected with an empty vector were used for control experiments. Other *A. rubens* neuropeptides [Luqin: EEKTRFPKFMRW-NH_2_ (ArLQ); tachykinin-like peptide 2: GGGVPHVFQSGGIFG-NH_2_ (ArTK2); (Semmens et al. 2016; Yañez-Guerra et al. 2018)] were tested at a concentration of 10 μM to assess the specificity of receptor activation.

### Generation and characterisation of antibodies to ArCCK1

To facilitate immunohistochemical analysis of the expression of a ArCCKP-derived neuropeptide in *A. rubens*, we generated antibodies to ArCCK1. To accomplish this an N-terminally truncated peptide analog of ArCCK1 with the addition of a reactive N-terminal lysine residue was synthesized (KY(SO_3_H)GHGLFW-NH_2,_ Peptide Protein Research Ltd, Fareham, UK). This peptide was conjugated to porcine thyroglobulin (Sigma-Aldrich, Gillingham, UK) as a carrier protein using 5% glutaraldehyde (Sigma-Aldrich, Gillingham, UK)in phosphate buffer (0.1 M; pH 7.2) and the conjugate was used for immunisation of a rabbit (70-day protocol; Charles River Biologics, Romans, France). The antigen was emulsified in Freund’s complete adjuvant for primary immunisations (∼100 nmol antigen peptide) and in Freund’s incomplete adjuvant for three booster immunisations (∼50 nmol antigen peptide). The presence of antibodies to the antigen peptide in post-immunisation serum samples was assessed using an enzyme-linked immunosorbent assay (ELISA), in comparison with pre-immune serum (Figure 5 – figure supplement 1a). Antibodies to the antigen peptide were purified from the final bleed antiserum by affinity-purification using the AminoLink Plus Immobilization Kit (ThermoFisher Scientific, Waltham, MA, USA), with bound antibodies eluted using glycine elution buffer [6.3 ml of 100 mM glycine (VWR Chemicals, Leicestershire, UK) and 0.7 ml of Tris (1M, pH = 7.0)] and trimethylamine (TEA) elution buffer [6.3 ml of TEA (Sigma-Aldrich, Gillingham, UK) and 0.7 ml of Tris (1M, pH = 7.0)]. Eluates were dialyzed and sodium azide (0.1%) was added for long-term storage of the affinity-purified antibodies at 4°C. The specificity of antibodies eluted with TEA, which were subsequently used for immunohistochemistry (see below), was assessed by ELISA by testing them at a concentration of 1:10 with the following synthetic peptides [100 µl at a concentration of 1 µM dissolved in carbonate/bicarbonate buffer (25 mM sodium carbonate, 25 mM sodium bicarbonate, pH = 9.8)]: ArCCK1, ArCCK2, ArCCK2(ns) and ArLQ (Figure 5 – figure supplement 1b). The rabbit antiserum to ArCCK1 has been assigned the RRID:AB_2877176.

### Immunohistochemical localisation of ArCCK1 in *A. rubens*

Small specimens of *A. rubens* (< 6 cm diameter) were fixed by immersion in seawater Bouin’s fluid [75% saturated picric acid (Sigma-Aldrich, Gillingham, UK) in seawater, 25% formaldehyde, 5% acetic acid] for 3 to 4 days at 4° C and then were decalcified for a week using a 2% ascorbic acid/0.3 M sodium chloride solution, dehydrated and embedded in paraffin wax. Sections of the arms and the central disk region (8 μm; transverse or horizontal) were cut using a microtome (RM 2145, Leica Microsystems [UK], Milton Keynes, UK) and mounted on chrome alum/gelatin coated microscope slides. Paraffin wax was removed by immersion of slides in xylene, and then slides were immersed in 100% ethanol. Endogenous peroxidase activity was quenched using a 0.3% hydrogen peroxide (VWT Chemicals, Leicestershire, UK)/methanol solution for 30 min. Subsequently, the slides were rehydrated through a graded ethanol series (90, 70 and 50%) and distilled water, blocked in 5% goat serum (NGS; Sigma-Aldrich, Gillingham, UK) made up in PBS containing 0.1% Tween (PBST). Then, the slides were incubated overnight with affinity-purified rabbit antibodies to ArCCK1 (TEA fraction diluted 1:10 in 5% NGS/PBST). Following a series of washes in PBST, indirect immunohistochemical detection was carried out using Peroxidase-AffiniPure Goat Anti-Rabbit IgG (H+L) conjugated to Horseradish Peroxidase (Jackson ImmunoResearch, West Grove, PA) diluted 1:1000 in 2% NGS/PBST. Bound antibodies were revealed using a solution containing 0.015% hydrogen peroxide, 0.05% diaminobenzidine (VWR Chemicals, Leicestershire, UK) and 0.05% nickel chloride (Sigma-Aldrich, Gillingham, UK) in PBS. When strong staining was observed, sections were washed in distilled water, dehydrated through a graded ethanol series (50, 70, 90 and 100%) and washed in xylene before being mounted with coverslips on DPX mounting medium (ThermoFisher Scientific, Waltham, MA, USA). Immunostaining was not observed in negative control tests without the primary antibodies or with primary antibodies that had been pre-adsorbed with the antigen peptide at a concentration of 20 µM (data not shown).

### Imaging

Photographs of sections processed for mRNA *in situ* hybridization or immunohistochemistry were captured using a QIClick^TM^ CCD Color Camera (Qimagin, British Columbia, CA) linked to a DMRA2 light microscope (Leica), utilising Volocity® v.6.3.1 image analysis software (PerkinElmer, Seer Green, UK) running on iMac computer (27-inch, Late 2013 model with OS X Yosemite, version 10.10). Montages of photographs were prepared using Adobe Photoshop CC (version 19.1.4, x64) running on a MacBook Pro computer (13-inch, early 2015 model with OS Mojave version 10.14.3).

### *In vitro* pharmacology

Informed by analysis of the expression of ArCCKP transcripts and ArCCK1 in *A. rubens*, both ArCCK1 and ArCCK2 were tested for myoactivity on cardiac stomach, tube foot, and apical muscle preparations dissected from specimens of *A. rubens* (n = 5 - 9, 8 - 10 and 20 - 23 respectively) and set up in a 20 ml organ bath, as described previously (Elphick et al. 1995, Melarange and Elphick 2003, Tian et al. 2017). Effects of peptides on preparations were assessed and recorded using an isotonic transducer (MLT0015, ADInstruments Pty Ltd) connected to a bridge amplifier (FE221 Bridge Amp, ADInstruments Pty Ltd) linked to PowerLab data adquisition hardware (2/36, ADInstruments Pty Ltd). Data was collected and analysed using LabChart (v8.0.7) software installed on a laptop computer (Lenovo E540, Windows 7 Professional). Stock solutions of synthetic peptides were prepared in distilled water and added to the organ bath to achieve final concentrations ranging from 0.1 nM to 1 μM. To assess the viability of preparations and to enable normalization of responses to ArCCK1 or ArCCK2, the starfish neuropeptide NGFFYamide (100 nM) was tested on cardiac stomach preparations and acetylcholine (ACh; 10 µM) was tested on tube foot and apical muscle preparations. To assess the importance of tyrosine sulphation for peptide bioactivity, a non-sulphated analog of ArCCK2 [ArCCK2(ns)] was also tested on cardiac stomach (n = 5), tube foot and apical muscle preparations (data not shown).

### *In vivo* pharmacology: testing ArCCK1 and ArCCK2 as cardiac stomach retractants

*In vitro* pharmacological experiments revealed that both ArCCK1 and ArCCK2 cause contraction of cardiac stomach preparations. Previous studies have revealed that the neuropeptide NGFFYamide causes cardiac stomach contraction *in vitro* and also triggers retraction of the everted cardiac stomach when injected into *A. rubens in vivo* (Semmens et al. 2013). Therefore, experiments were performed to investigate if ArCCK1 and ArCCK2 also trigger cardiac stomach retraction in *A. rubens*. Twenty specimens of *A. rubens*, which had been withheld from a food supply for one week, were placed in a glass tank containing 2% magnesium chloride (MgCl_2_; Sigma-Aldrich, Gillingham, UK) dissolved in seawater, which acts as a muscle relaxant in marine invertebrates (Mayer, 1909). This treatment conveniently and reproducibly causes eversion of the cardiac stomach in *A. rubens*, typically within a period of ∼30 min (Semmens et al. 2013). Hamilton 75N 10 μl syringes (Sigma-Aldrich, Gillingham, UK) were used to inject test compounds into the perivisceral coelom of animals, inserting the needle through the aboral body wall of the arms proximal to the junctions with the central disk region. Care was taken to inject neuropeptides [ArCCK1, ArCCK2 and ArCCK2(ns)] or distilled water (control) into the perivisceral coelom and not into the cardiac stomach. All animals were first injected with 10 μl of distilled water (control) and video recorded for 6 min. The same animals were then injected with 10 μl of 1 mM peptide and video recorded for 6 min. Static images from video recordings were captured at 30 s intervals from the time of injection. Then the two-dimensional area of everted cardiac stomach was measured from the images using the ImageJ software (version 1.0; http://rsb.info.nih.gov/ij) and normalized as a percentage of the area of cardiac stomach everted at the time of injection (T0).

### *In vivo* pharmacology: testing effects of ArCCK1 and ArCCK2 on feeding behaviour

Previous studies have revealed that the starfish neuropeptide NGFFYamide inhibits the onset of feeding behaviour of *A. rubens* on a prey species – the mussel *Mytilus edulis* (Tinoco et al. 2018). Here the same methods employed by Tinoco et al. (2018) were used to investigate if ArCCK1 and/or ArCCK2 affect feeding behaviour in starfish. Sixty-two adult starfish (n = 24 for ArCCK1; n = 38 for ArCCK2) that met the following criteria were used: *(i)* all 5 arms were intact, *(ii)* exhibited a normal righting response (Lawrence and Cowell 1996) and *(iii)* after twenty-four days of starvation, exhibited normal feeding behaviour on a mussel. Then, starfish were fasted for twenty-four days and transferred to and kept individually in Plexiglas aquaria, as described previously (Tinoco et al. 2018). After three days of acclimation (twenty-seven days of starvation at this point) and at 10 a.m., these animals were then divided into a control group (to be injected with distilled water), with 13 animals used for the ArCCK1 experiment (mean diameter of 12.4 ± 0.3 cm) and 19 animals used for the ArCCK2 experiment (mean diameter of 12.9 ± 0.4 cm), and a test group (to be injected with ArCCK1 or ArCCK2), with 11 animals used for the ArCCK1 experiment (mean diameter of 12.6 ± 0.3 cm) and 19 animals used for the ArCCK2 experiment (mean diameter of 12.9 ± 0.5 cm). The starfish were then injected with 10 µl of distilled water (control group) or 10 µl of 1 mM ArCCK1 or ArCCK2 peptides (test group) to achieve an estimated final concentration in the perivisceral coelom of ∼1 µM, which is the concentration at which ArCCK peptides were found to have a maximal effect when tested on *in vitro* preparations of the cardiac stomach. The time taken for starfish to make first contact with a mussel (tube feet touching the mussel or time to touch the mussel), the number of attempts to touch as well as the time to enclose the mussel (indicated by a feeding posture) was recorded. Starfish that were feeding after 24 h were included in data analysis and any starfish in the control or test group that had not fed on a mussel after 24 hours were discarded from data analysis.

### Statistical analyses

Data were presented as means ± standard error of the mean (s.e.m). The *in vitro* or *in vivo* pharmacological effects of starfish CCK-type peptides on cardiac stomach, apical muscle and tube foot preparations were analysed by 2-way ANOVA, using type of substance tested and concentration/time as independent factors and Bonferroni’s multiple comparison test. Apical muscle and tube foot data were transformed to logarithms to obtain a normal distribution and homogeneity of variances. The *in vitro* effects of ArCCK1 and ArCCK2 (1 µM) on cardiac stomach preparations were compared with the *in vitro* effect of NGFFYamide (100 nM) using a two-tailed Student’s t-test. The effect of ArCCK1 on feeding behaviour was analysed by two-tailed Mann-Whitney U-test (time to touch and time to enclose) because these data did not follow a normal distribution when analysed using the D’Agostino & Pearson omnibus normality test. The effect of ArCCK2 on feeding behaviour was analysed by two-tailed Student’s t-test (time to touch) or Welch’s unequal variances t-test (time to enclose). Fisher’s exact test was used to analyse the percentage of successful feeding after the first touch for control and treated starfish. Statistical analyses were carried out using Prism 6 (GraphPad software, La Jolla, CA, USA) and differences were considered statistically significant at p < 0.05.

## Acknowledgments

This work was supported by grants awarded to MRE by BBSRC (BB/M001644/1) and the Leverhulme Trust (RPG-2018-200) and to AMJ by BBSRC (BB/M001032/1). LAYG was supported by a PhD studentship awarded by the Mexican Council of Science and Technology (CONACyT studentship no. 518612) and Queen Mary University of London. YZ was supported by a PhD studentship awarded by the China Scholarship Council and Queen Mary University of London. JD is currently a postdoctoral researcher supported by Fund for Scientific Research of Belgium (F.R.S.-FNRS).

## Supplementary data

**Figure 1 – source data 1.**
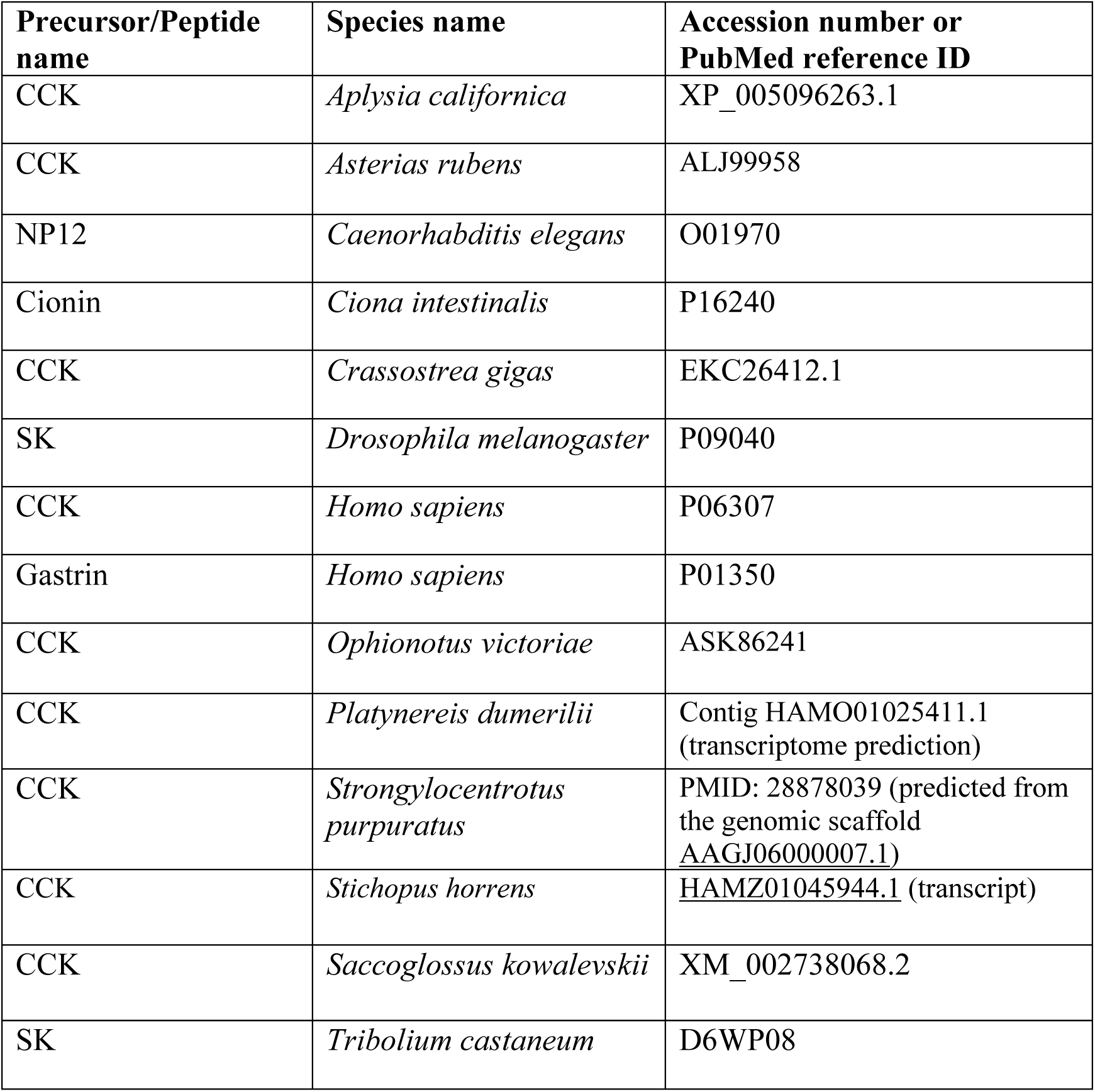
Accession numbers for precursors of the neuropeptides shown in the sequence alignment in Figure 1.

**Figure 1 – source data 2.** Data for the mass spectra shown in Figure 1 – figure supplement 2. (not included here – available as separate file)

**Figure 1 – figure supplement 1.**
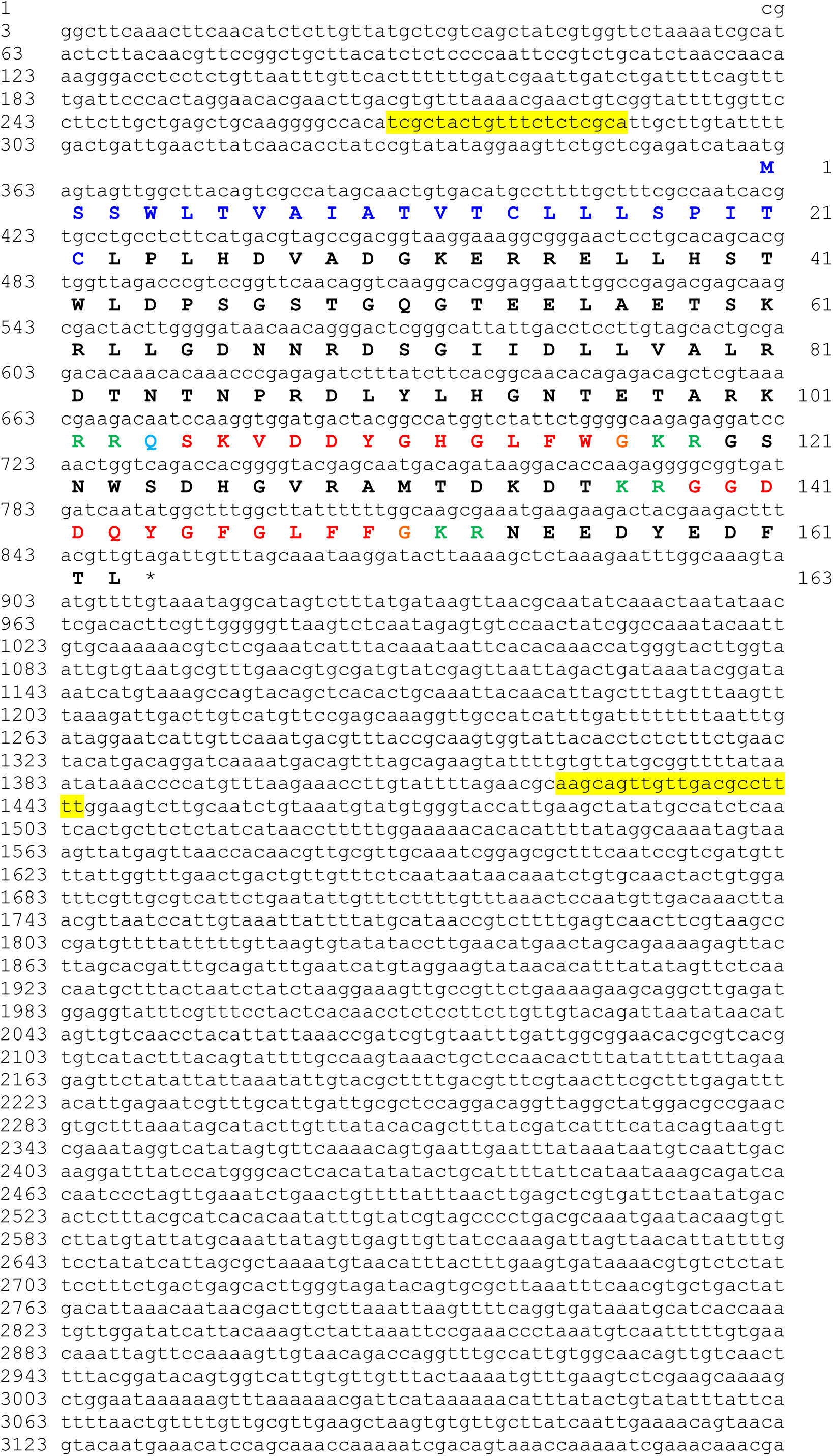
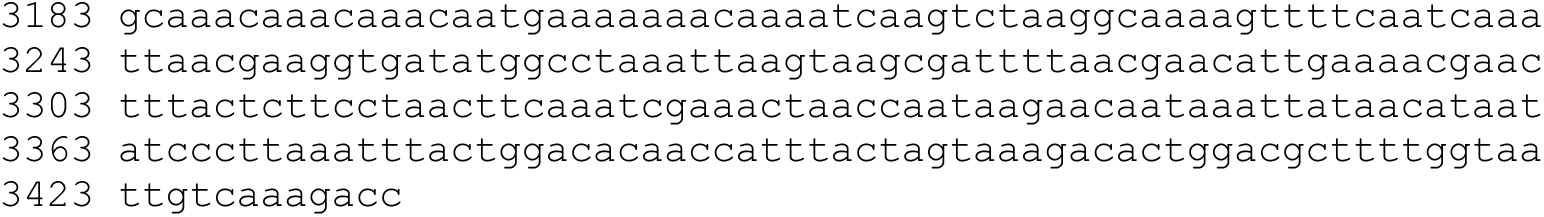
The *A. rubens* CCK-type precursor (ArCCKP). The nucleotide sequence of contig 1124413 derived from radial nerve cord transcriptome data (Semmens et al., 2016; GenBank accession number KT601716) is shown in lowercase (3434 bases) and the encoded the precursor protein sequence is shown in uppercase (163 residues). The predicted signal peptide is shown in dark-blue, two putative CCK-type peptides are shown in red but with an N-terminal glutamine residue in first peptide shown light blue to indicate that it is a potential substrate for pyroglutamination. Putative dibasic cleavage sites are shown in green. The asterisk shows the position of the stop codon. The sequences of primers used for PCR cloning of a cDNA encoding ArCCKP are highlighted in yellow. The sequence of the cloned cDNA was found to be identical to the corresponding sequence of contig 1124413.

**Figure 1 – figure supplement 2.**
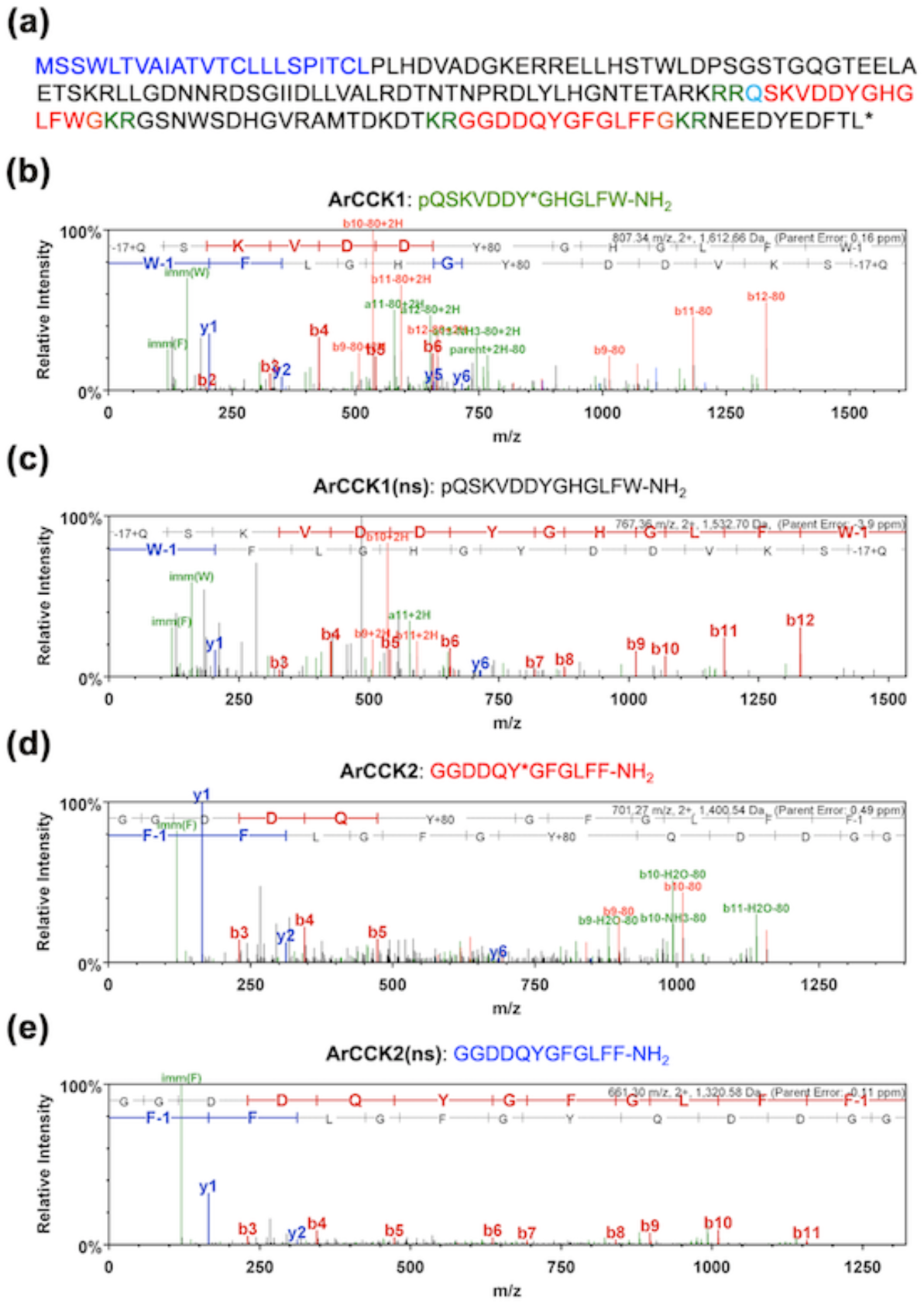
Determination of the structures of peptides derived from the ArCCKP by mass spectrometric (LC-MS-MS) analysis of *A. rubens* radial nerve cord extract. **(a)** Amino acid sequence of ArCCKP, with the predicted signal peptide shown in dark blue, predicted dibasic cleavage sites shown in green and predicted CCK-type neuropeptides shown in red. C-terminal glycine residues that are predicted substrates for amidation are shown in orange and an N-terminal glutamine residue that is a potential substrate for pyroglutamination (pQ) is shown in light blue. The nucleotide sequence of a cDNA encoding ArCCKP is shown in Figure 1 – figure supplement 1. **(b – d)** MS/MS data for four CCK-type peptides derived from ArCCKP detected in *A. rubens* radial nerve cord extracts: **(b)** pQSKVDDY(SO_3_H)GHGLFW-NH_2_ [ArCCK1, which has a sulphated tyrosine], **(c)** pQSKVDDYGHGLFW-NH_2_ [ArCCK1(ns), which is not sulphated], **(d)** GGDDQY(SO_3_H)GFGLFF-NH_2_ [ArCCK2, which has a sulphated tyrosine] and **(e)** GGDDQYGFGLFF-NH_2_ [ArCCK2(ns), which is not sulphated]. The b series of peptide fragment ions are shown in red, the y series are shown in blue and additional identified peptide fragment ions are shown in green. The amino acid sequence identified in each mass spectrum is shown above it, with −17+Q representing an N-terminal pyroglutamate residue (pQ), W-1 representing an amidated C-terminal tryptophan residue (W-NH_2_), F-1 representing an amidated C-terminal phenylalanine residue (F-NH_2_) and Y+80 representing a sulphated tyrosine residue (Y*). The observed m/z of the precursor ion for each peptide is with a charge state of +2 and with errors between the experimentally determined and predicted values ranging from −3.9 ppm to +0.49 ppm.

**Figure 2 – source data 1.**
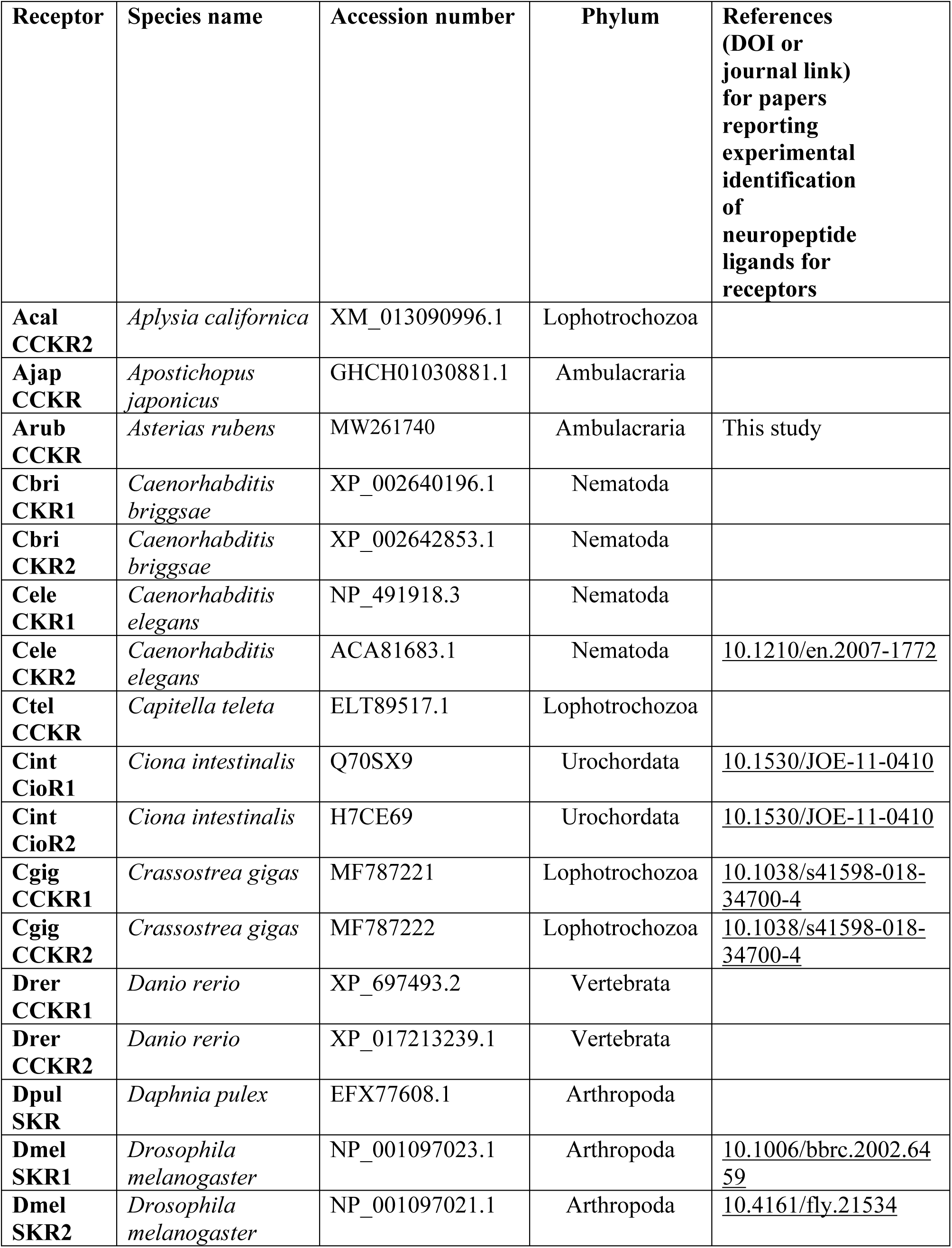
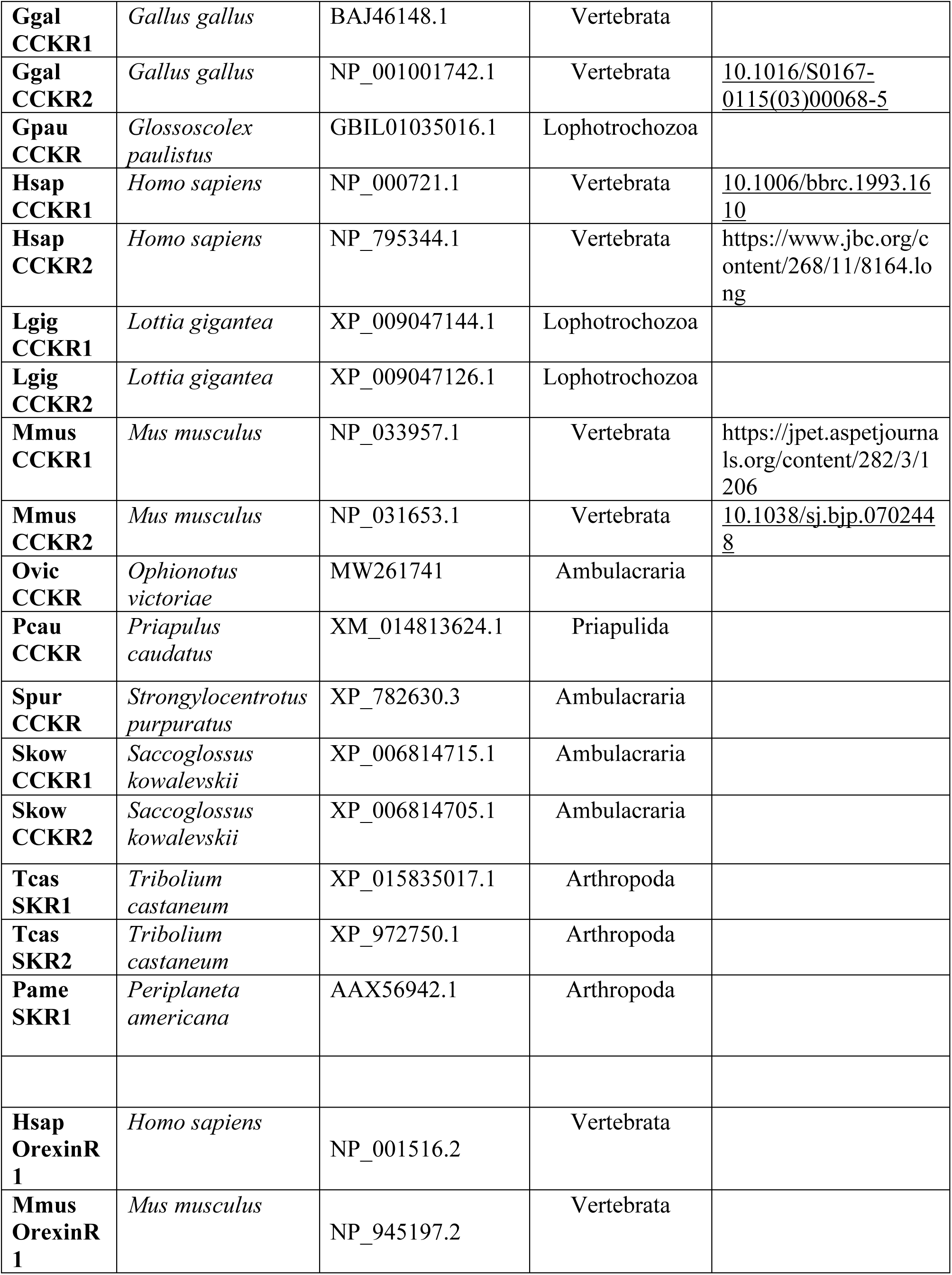
Accession numbers for the receptor sequences used for the phylogenetic tree in Figure 2.

**Figure 2 - figure supplement 1.**
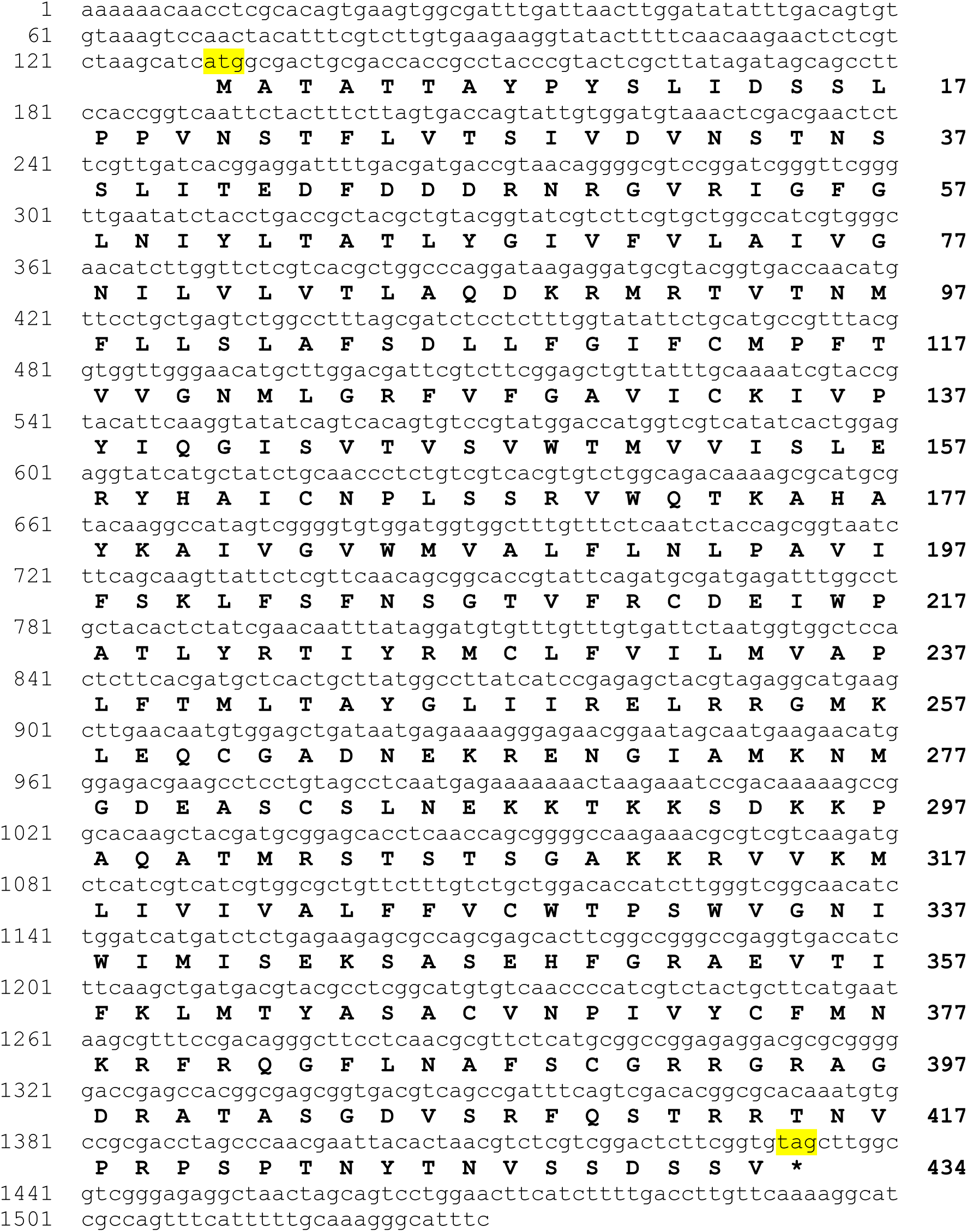
The *A. rubens* CCK receptor (ArCCKR). The nucleotide sequence of contig 1110296 derived from radial nerve cord transcriptome data (GenBank accession number MW261740) is shown in lowercase (1530 bases; numbering on the left) and the encoded ArCCKR protein sequence is shown in uppercase (434 residues). The start and stop codons are highlighted in yellow. The coding sequence was synthesized (GenScript) to enable testing of the ArCCKP-derived CCK-type peptides as ligands for ArCCKR (see Figure 3).

**Figure 2 – figure supplement 2.**
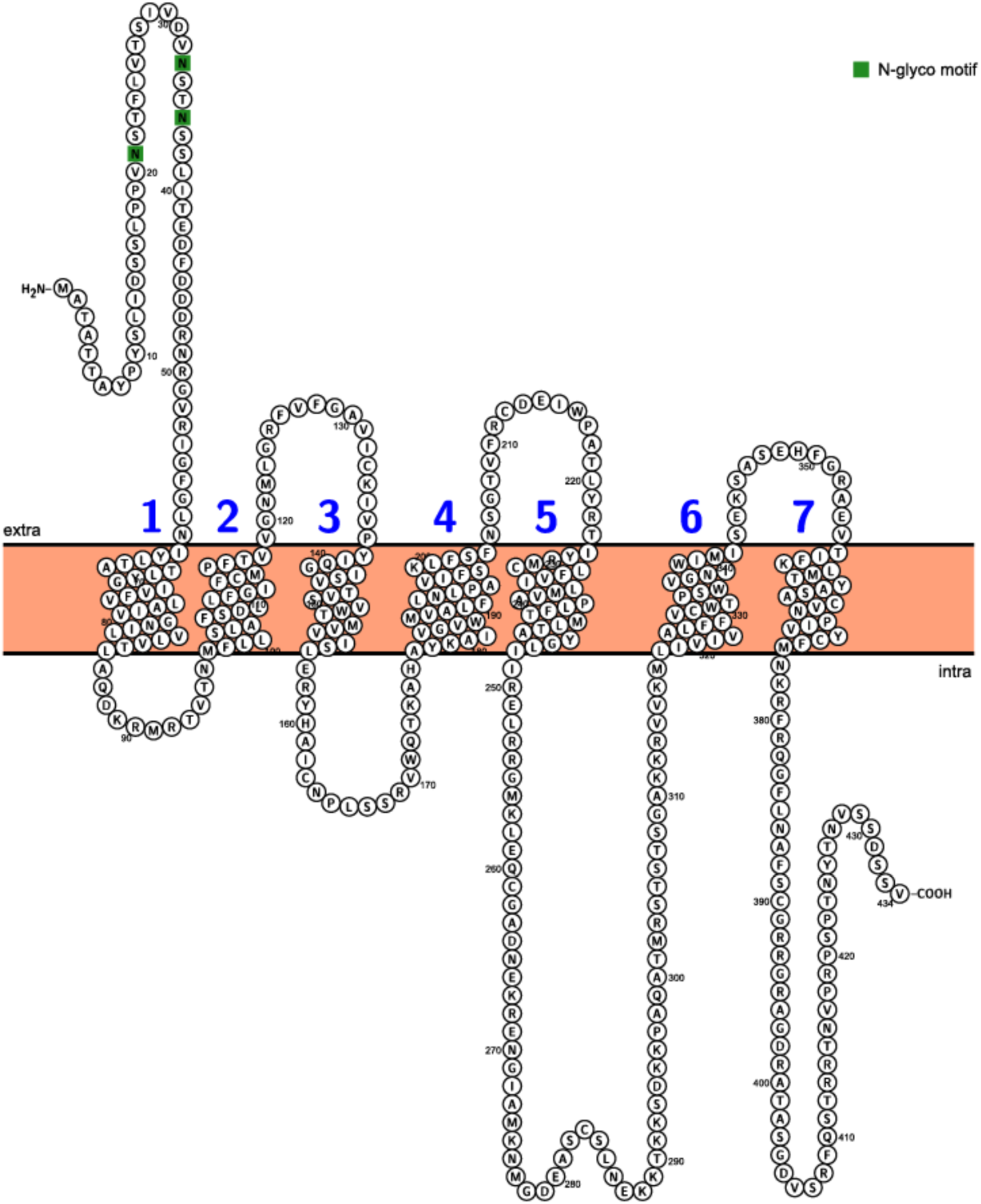
Topology of ArCCKR. Predicted topology of ArCCKR inferred by the Protter tool (Omasits et al. 2014) with seven transmembrane domains numbered in blue and predicted N-glycosylation sites shown in green.

**Figure 3 – source data 1.** Data for the graphs shown in Figure 3 and Figure 3 – figure supplement 1. (not included here – available as separate file)

**Figure 3 – figure supplement 1.**
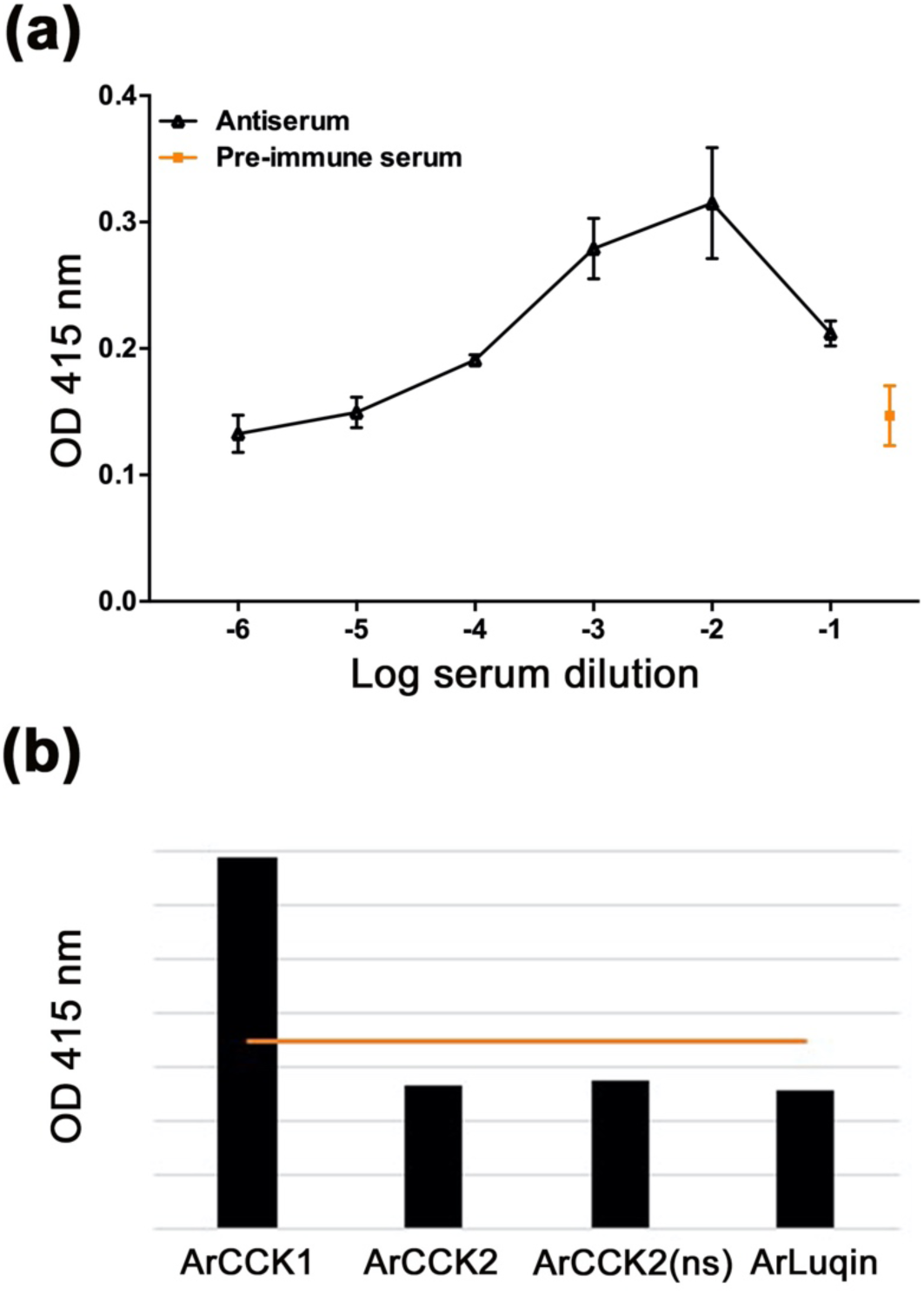
Graph showing the selectivity of ArCCKR as a receptor for CCK-type peptides. The *A. rubens* CCK-type peptide ArCCK2 (red) triggers luminescence in CHO-K1 cells expressing the receptor ArCCKR, the promiscuous G-protein Gα16 and the calcium-sensitive luminescent GFP-apoaequorin fusion protein G5A. In these experiments ArCCK2 was tested at concentrations between 10^-8^ and 10^-4^ M; see Figure 3b for experiments showing complete concentration-response curves. The receptor is not activated by the SALMFamide-type neuropeptide S2 (SGPYSFNSGLTF-NH_2_; purple) or by the *A. rubens* tachykinin-like peptide ArGxFamide2 (GGGVPHVFQSGGIF-NH_2_; orange). Each point represents mean values (± s.e.m) from at least three independent experiments done in triplicate.

**Figure 5 – figure supplement 1.**
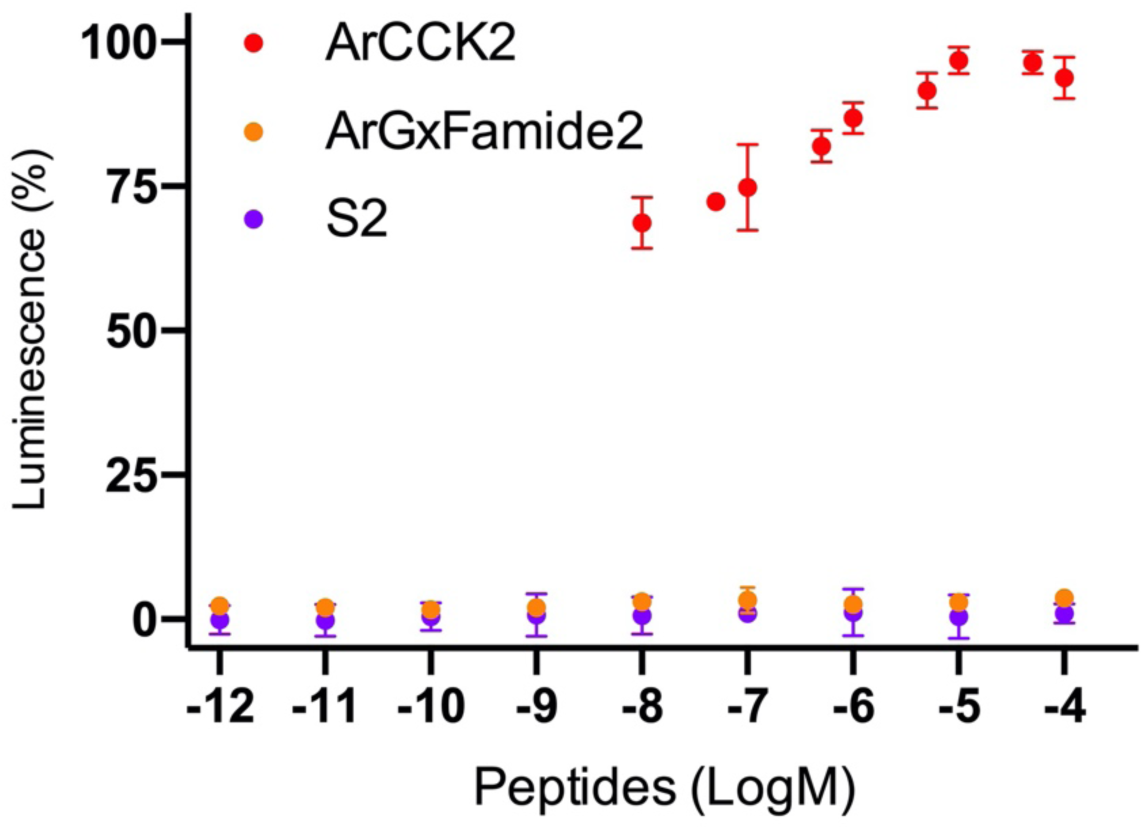
Characterisation of a rabbit antibodies to ArCCK1 using an enzyme-linked immunosorbent assay (ELISA) **(a)**. By comparison with pre-immune (orange), 0.1 nmol of the ArCCK1 antigen peptide is detected by the ArCCK1 antiserum (black line) at dilutions between 1:10 and 1:10000. All data points are mean values from two replicates. **(b)** Graph showing the results of ELISA tests using the TEA fraction of affinity-purified antibodies to ArCCK1 (dilution 1:10). The red line indicates the mean optical density (OD) value of negative control experiments without peptides. The four peptides tested were applied at a concentration of 0.1 nmol but only the mean OD value for ArCCK1 is above the OD value for the negative control. All data points are mean values from six replicates. These experiments demonstrate that specific antibodies to ArCCK1 were successfully generated.

**Figure 6 – source data 1.** Data for graphs shown in Figure 6b, d and f. (not included here – available as separate file)

**Figure 7 – source data 1.** Data for graphs shown in Figure 7a, b and c. (not included here – available as separate file)

**Figure 7 - video 1. ArCCK1** (10 µl 1 mM) **induced retraction of the cardiac stomach in the starfish *A. rubens***. (not included here – available as separate file)

**Figure 7 - video 2. ArCCK2** (10 µl 1 mM) **induced retraction of the cardiac stomach in the starfish *A. rubens***. (not included here – available as separate file)

**Figure 8 – source data 1.** Data for graphs shown in Figure 8. (not included here – available as separate file)

## Key Resources Table

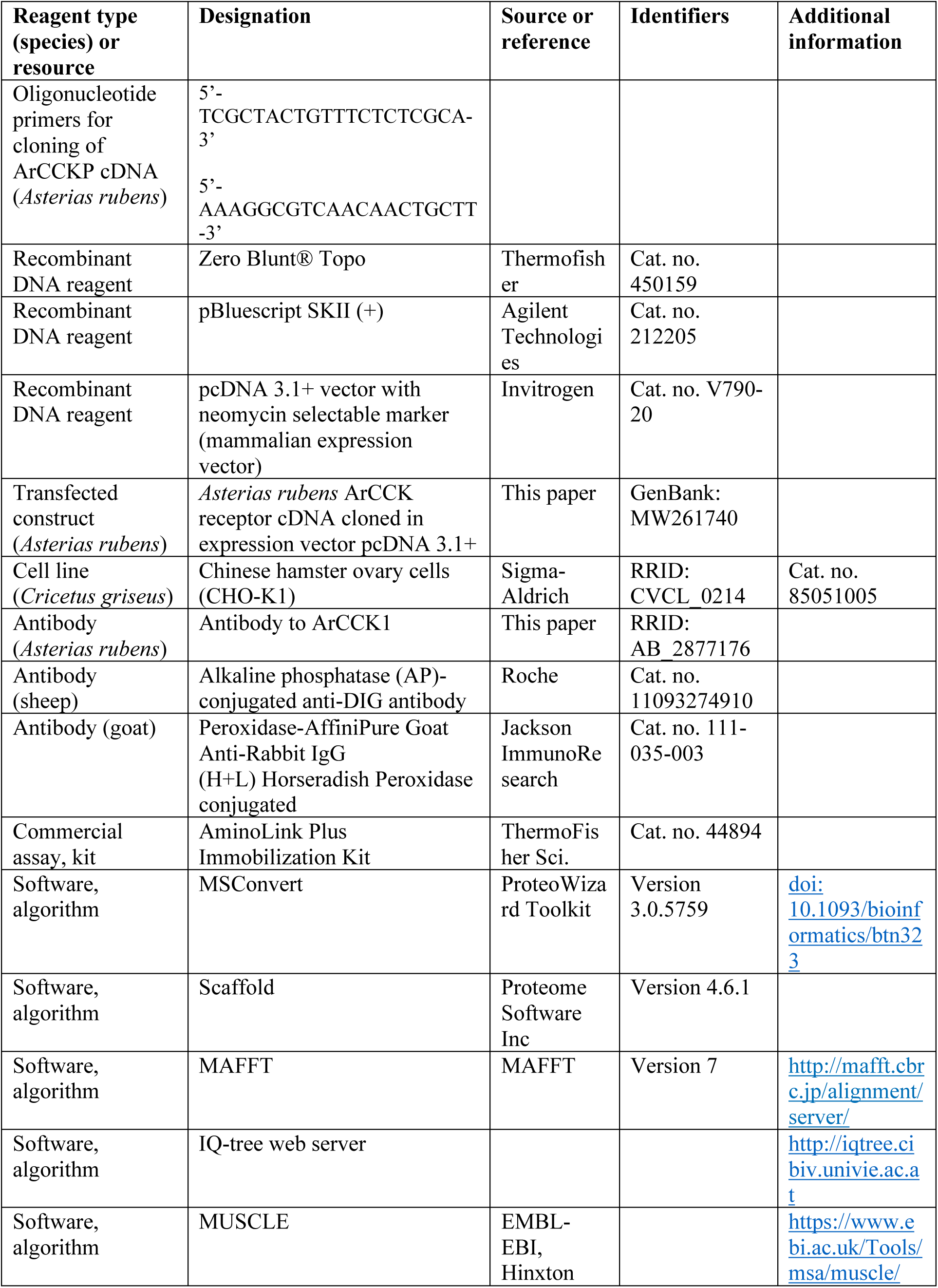
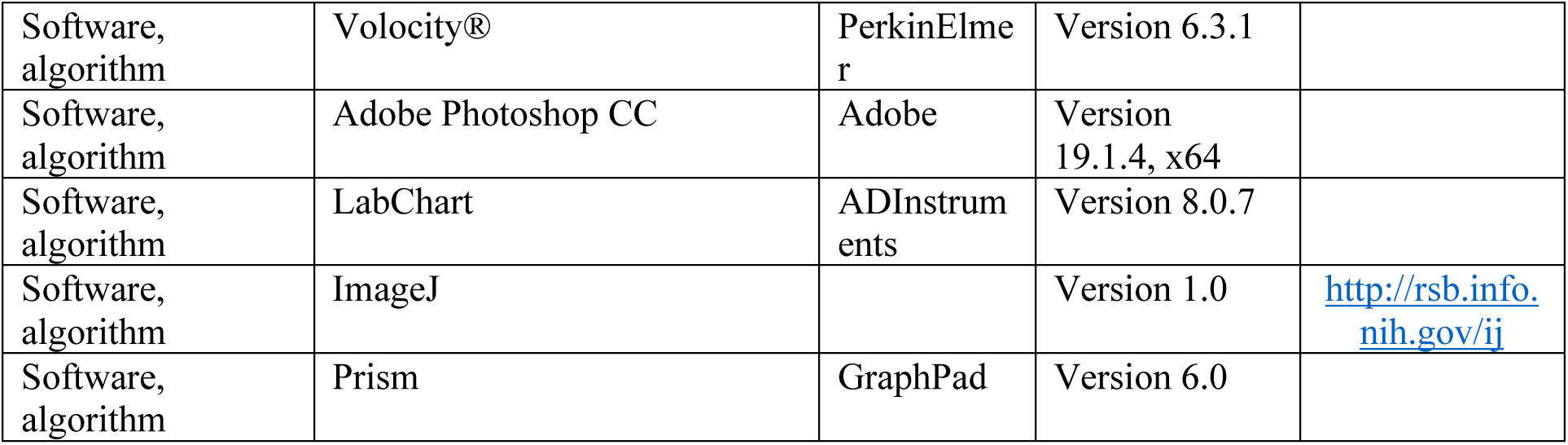

## Notes

### Competing Interest Statement

The authors have declared no competing interest.

